# Dynamics of social representation in the mouse prefrontal cortex

**DOI:** 10.1101/321182

**Authors:** Dana Rubi Levy, Tal Tamir, Maya Kaufman, Aharon Weissbrod, Elad Schneidman, Ofer Yizhar

## Abstract

The prefrontal cortex (PFC) plays an important role in regulating social functions in mammals, and impairments in this region have been linked with social dysfunction in psychiatric disorders. Yet little is known of how the PFC encodes social information and of how social representations may be altered in such disorders. Here, we show that neurons in the medial PFC (mPFC) of freely behaving mice preferentially respond to socially-relevant sensory cues. Population activity patterns in the mPFC differed considerably between social and nonsocial stimuli and underwent experience-dependent refinement. In Cntnap2 knockout mice, a genetic model of autism, both the categorization of sensory stimuli and the refinement of social representations were impaired. Noise levels in spontaneous population activity were higher in Cntnap2 mice, and correlated strongly with the degree to which social representations were disrupted. Our findings elucidate the encoding of social sensory cues in the mPFC, and provide an important link between altered prefrontal dynamics and autism-associated social dysfunction.

## Introduction

Social interactions are a central aspect of animal behavior, and are orchestrated by multiple neural circuits throughout the brain^1^. The complexity of social behaviors requires constant integration of sensory cues^2,3^ with internal motivational and arousal states^1,4^, and the coordination of intricate motor sequences^5–7^. The prefrontal cortex (PFC) is known to integrate such internal and external variables^8^, and is crucial for social functions in humans^9,10^ and other animals^11^–16. Neurons in the PFC represent multiple aspects of the external world, responding to salient sensory cues associated with positive^17^ or negative^18^ reinforcement, and display mixed selectivity to combinations of task-related variables^19,20^. In social contexts, neural activity in the PFC was shown to increase during approach toward a conspecific^12,15^ and represent both spatial and social aspects of behavior^14^. Yet, little is known regarding the response selectivity of PFC neurons to social sensory cues, and the dynamics of social representations in prefrontal circuits remains largely unexplored.

Impairments of the PFC have been widely reported in autism spectrum disorder (ASD)^21–26^, a neurodevelopmental disorder associated with altered social function. Imaging studies have identified reduced long-range prefrontal connectivity in individuals with ASD^27–29^, and demonstrated poor selectivity to sensory stimuli^30^, as well as higher trial-to-trial variability in multiple cortical sensory areas^29,31^. These findings are consistent with a prominent theory of ASD pathophysiology, which suggested that ASD arises from developmental changes in the balance of neocortical excitation and inhibition (E/I balance)^32^. This cortical imbalance was hypothesized to disrupt cortical maturation and long-range connectivity, resulting in prominent changes in information processing and elevated cortical noise^32–34^. Work in animal models provided additional support to this hypothesis^35–37^, despite conflicting findings regarding the nature of E/I imbalance^36,38,39^. Yet, most of our knowledge regarding autism-associated changes in neuronal functional properties is based on ex-vivo studies, and despite extensive behavioral and molecular phenotyping of mouse models of autism, not much is known about the emergent changes in circuit dynamics in vivo^36,39–44^.

We studied the representation of social information in the medial prefrontal cortex (mPFC) of freely-behaving mice. To characterize the nature of neural coding and stimulus processing in social dysfunction, we compared neural activity in wild-type (wt) and Cntnap2 knockout (Cntnap2^−/−^) mice, an established genetic model of autism^45^. We found that in wild-type mice, mPFC neurons 3 displayed robust response selectivity to social over nonsocial sensory cues. Population-level analysis revealed distinct categorization of sensory cues based on their social nature, which underwent marked experience-dependent refinement over experimental sessions. In Cntnap2^−/−^ mice, mPFC activity showed reduced differentiation between social and nonsocial stimuli and lacked experience-dependent dynamics. Strikingly, the deficits in social-specific activity patterns in Cntnap2^−/−^ mice were strongly correlated with elevated variability of spontaneous neuronal activity. Our results uncover distinct coding of social sensory cues in the mPFC and provide a potential link between mPFC E/I imbalance, altered encoding of socially-relevant stimuli and autism-associated social dysfunction.

## Results

### Medial prefrontal cortex neurons are tuned to social cues

Social behaviors in rodents are primarily guided by the emission and detection of specific chemosensory cues^4,46,47^. To study the responses of prefrontal neurons to social cues, we used a custom-built odor delivery system, which allowed for highly precise presentation of olfactory stimuli while recording mPFC activity in freely-behaving male mice (Fig. 1a,b; Supplementary Fig. 1). Each mouse was repeatedly presented (in pseudorandomized order) with the odors of male mice (M), female mice (F), and with three nonsocial odors of distinct valence: banana extract (B), considered to be a neutral stimulus to mice^48^; peanut oil (P), an attractive stimulus^48^; and hexanal (H), known to be mildly repellent^49^. In interleaved control trials, clean air (CA) was presented using the same odor delivery system. Mice displayed pronounced behavioral responses following presentation of the odors, which consisted of orienting toward the odor port and increased locomotion. The probability of odor-directed orientation responses was higher during delivery of social cues (Fig. 1c). However, the response latency (Fig. 1d) and the stimulus-evoked increase in locomotion (Fig. 1e) did not significantly differ between social and nonsocial stimuli.

**Figure 1.**
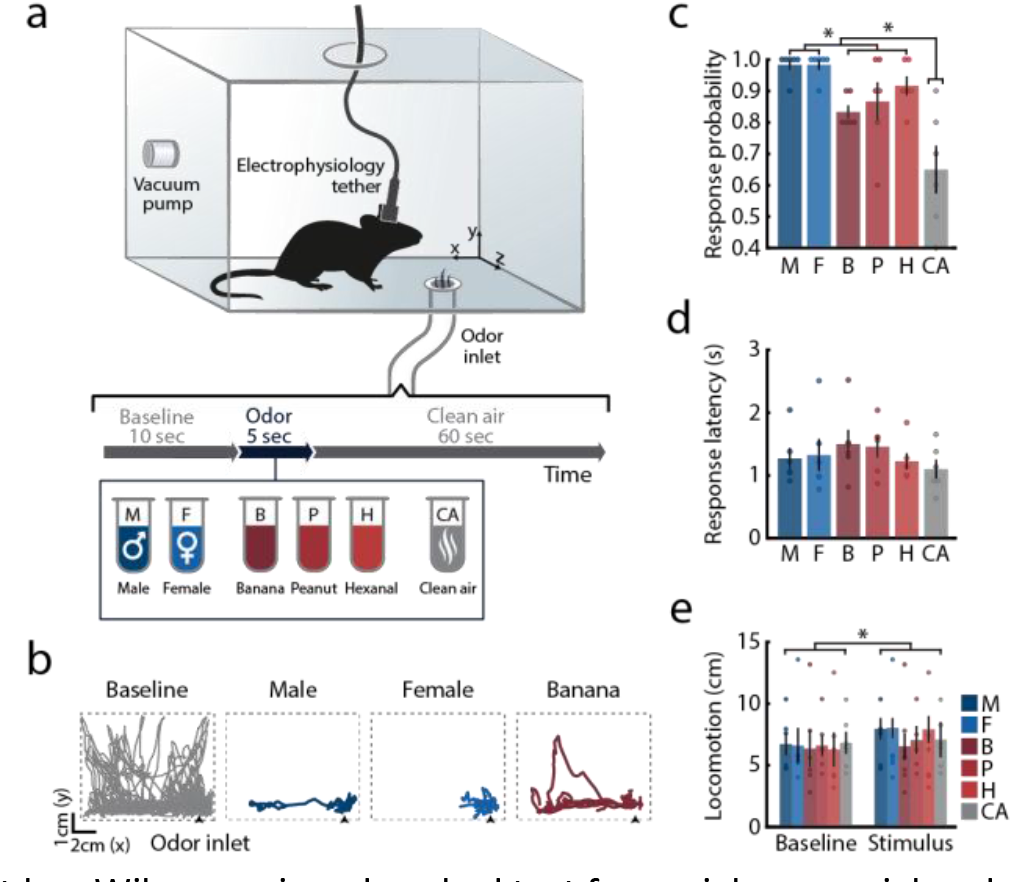
Experimental design: presentation of social and nonsocial sensory cues in freely-behaving mice. **(a)** Schematic representation of the experimental chamber and trial design: freely behaving male mice were presented with social and nonsocial olfactory cues. Odors were presented in pseudo-random order, interleaved with control trials where only clean air was presented. All trials were preceded and followed by constant infusion of clean air. Electrophysiological data was continuously recorded from the mPFC. **(b)** Representative side-view trajectories of mouse locomotion during pre-stimulus baseline periods (grey) and during stimulus presentations (color). **(c)** Probability of odor-evoked orientation responses across all odors. Friedman test for comparison of all stimuli, *χ*^2^_(5)_ = 21.235, *P* < 0.001; Friedman test with post hoc Wilcoxon signed-ranked test for social, nonsocial and clean air comparison, *χ*^2^_(2)_ = 12.0, *P* < 0.01, statistical significance of post hoc analysis is marked on figure (mean response probability: social = 98.33 ± 1.05, nonsocial = 87.22 ± 3.03, clean air = 65 ± 7.63). **(d)** Mean latency to odor-evoked orientation responses. Friedman test *χ*^2^_(5)_ = 1.809, *P* = 0.874. **(e)** Locomotion during 5 s of stimulus presentation and during the corresponding pre-stimulus baseline periods. Two-way repeated measures (RM) ANOVA, *F*phase(1,5) = 8.151, *P* < 0.05; *F*stimulus(5,25) = 0.387, *P* = 0.853; *F*phase*stimulus(5,25) = 0.822, *P* = 0.546. Color code represents stimulus identity; circles mark individual mice. Data is presented as mean ± SEM. *n* = 6, \**P* < 0.05.

We recorded stimulus-evoked responses of mPFC units (*n* = 194; 6 mice) and found that 44% responded to at least one stimulus, typically by increasing their firing rates (Fig. 2a and Supplementary Fig. 2a-c). Presentation of both male and female cues recruited more mPFC units than any of the nonsocial odors (Fig. 2b). Almost a quarter of recorded units displayed selectivity to social signals – twice the number of units with nonsocial odor selectivity or mixed social/nonsocial responses (Fig. 2c left and Supplementary Fig. 2c-e). In addition, the magnitude of neuronal responses to social odors was significantly higher (Fig. 2b,d and Supplementary Fig. 2f). Among responsive units, 51% were stimulus-specific, of which a large fraction responded exclusively to male or female cues (Fig. 2c, right). The average unit tuning, calculated as the normalized odor-evoked response magnitude in all units that showed stimulus-associated responses, was higher for both male and female cues than for all nonsocial odors, regardless of odor identity (Fig. 2e).

**Figure 2.**
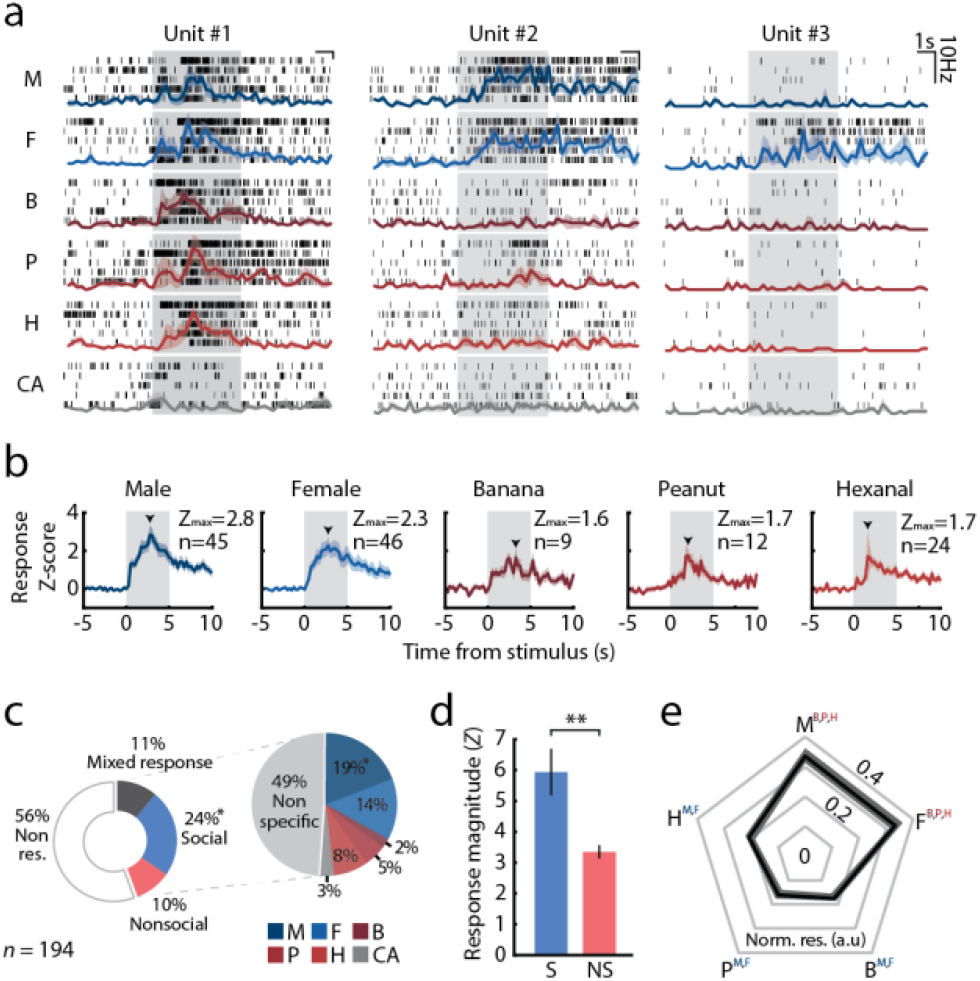
Distinct representation of social cues in the mPFC neuronal population code. **(a)** Schematic illustration of the population analysis. *Top*: multiple units recorded during a single recording session in response to each of the presented stimuli. Response patterns were used for neural trajectory analysis (as binned spike counts; panel b), and binarized in finer time bins for modeling the probability distribution of response patterns and response dissimilarity (*bottom*; panels c-e). **(b)** Representative 2D projections of the neuronal population trajectories before, during, and after stimulus presentation (each trajectory spans 5s, where each point was estimated in 150 ms bins, see Online Methods). The schemes above the panels indicate the corresponding stage along the trial. Colors represent odor identity. Here, the first two principal components accounted for 75% of the variance. **(c)** Similarity matrix depicting the population-based representation distance between each pair of stimuli, calculated over the final 2.5 s window of stimulus presentation. The block diagonal structure of the matrix indicates a clear divergence between social and nonsocial categories in the recorded mPFC activity. **(d)** Distances between population responses to male cues and all stimuli (including the “self-distance” between responses to male odor on different trials). Circles depict individual mice (*n* = 6 mice, 2 recording sessions for each mouse). RM ANOVA with Bonferroni corrected post hoc comparisons. *F*stimulus(4,40) = 16.255, *P* < 0.001. **(e)** Time-dependent distance of all stimulus representations from male cue responses, calculated in consecutive 1 s windows. Shaded area represents cue delivery time. Two-way RM ANOVA (for M, F and nonsocial stimuli) was performed with Dunnett’s comparisons for each stimulus against its last baseline bin. *F*stimulus*time(40,400) = 3.8397, *P* < 0.001; *F*stimulus (2,20) = 29.7942, *P* < 0.001; *F*_time (20,200)_ = 5.665, *P* < 0.001. Arrowheads mark range of post hoc statistical significance for nonsocial stimuli. For all panels: \*\*\**P* < 0.001, Mean ± SEM (shaded area/error bars) is presented. M, male; F, female; B, banana; P, peanut; H, hexanal; CA, clean air.

### Distinct population representations of social and nonsocial stimuli in the mPFC

To further elucidate the nature of prefrontal representation of social signals, we analyzed the activity patterns of simultaneously-recorded mPFC neurons (14–23 units per mouse; Fig. 3a). First, we discretized neural responses into 150 ms bins, and used Principal Component Analysis (PCA) to project the population firing rates (FR) as a function of time, onto the first two principal components (see Online Methods). This analysis revealed a clear divergence of population responses evoked by the social and nonsocial stimuli soon after stimulus onset (Fig. 3b). This category-specific separation of population trajectories persisted for several seconds after stimulus offset before converging back to baseline activity state (see Fig. 3b, right panel).

**Figure 3.**
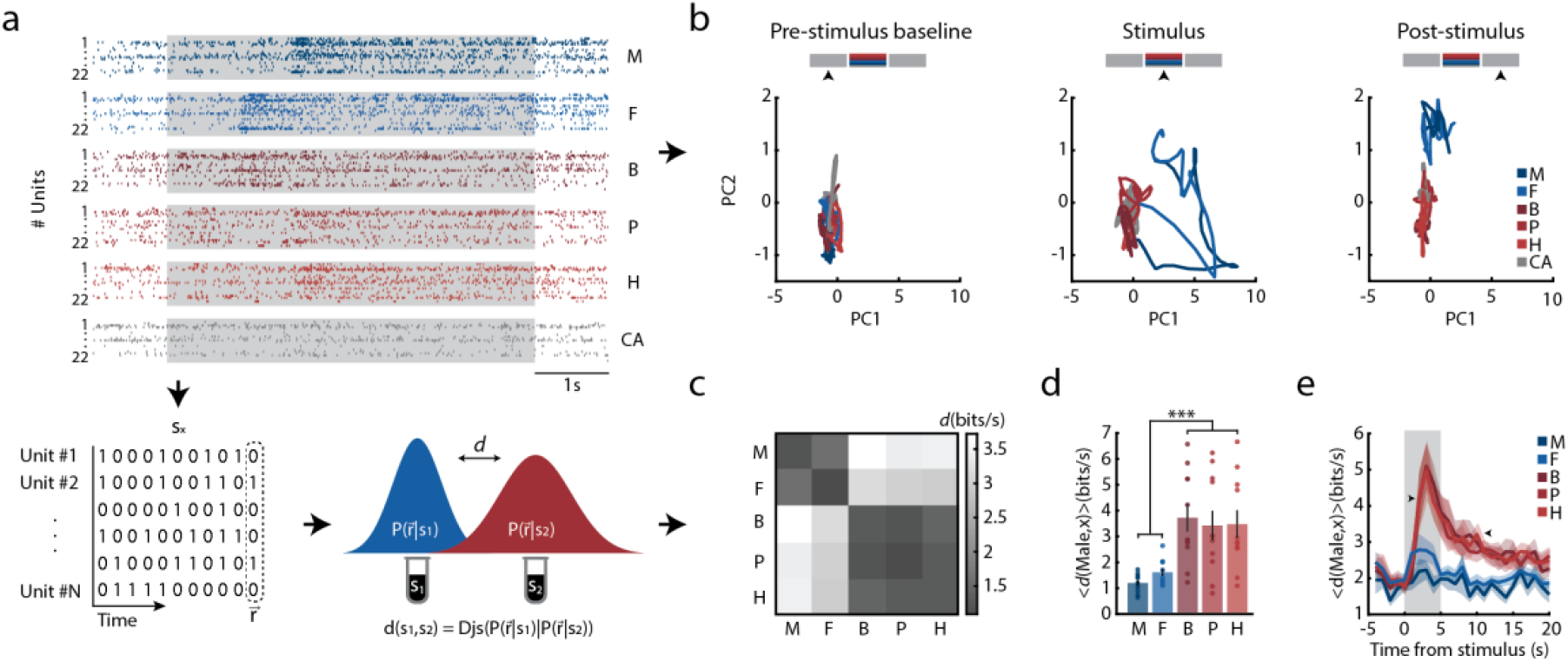
Distinct representation of social cues in the mPFC neuronal population code. **(a)** Schematic illustration of the population analysis. *Top*: multiple units recorded during a single recording session in response to each of the presented stimuli. Response patterns were used for neural trajectory analysis (as binned spike counts; panel b), and binarized in finer time bins for modeling the probability distribution of response patterns and response dissimilarity (*bottom*; panels c-e). **(b)** Representative 2D projections of the neuronal population trajectories before, during, and after stimulus presentation (each trajectory spans 5s, where each point was estimated in 150 ms bins, see Online Methods). The schemes above the panels indicate the corresponding stage along the trial. Colors represent odor identity. Here, the first two principal components accounted for 75% of the variance. **(c)** Similarity matrix depicting the population-based representation distance between each pair of stimuli, calculated over the final 2.5 s window of stimulus presentation. The block diagonal structure of the matrix indicates a clear divergence between social and nonsocial categories in the recorded mPFC activity. **(d)** Distances between population responses to male cues and all stimuli (including the “self-distance” between responses to male odor on different trials). Circles depict individual mice (*n* = 6 mice, 2 recording sessions for each mouse). RM ANOVA with Bonferroni corrected post hoc comparisons. *F*stimulus(4,40) = 16.255, *P* < 0.001. **(e)** Time-dependent distance of all stimulus representations from male cue responses, calculated in consecutive 1 s windows. Shaded area represents cue delivery time. Two-way RM ANOVA (for M, F and nonsocial stimuli) was performed with Dunnett’s comparisons for each stimulus against its last baseline bin. *F*stimulus*time(40,400) = 3.8397, *P* < 0.001; *F*stimulus (2,20) = 29.7942, *P* < 0.001; *F*_time (20,200)_ = 5.665, *P* < 0.001. Arrowheads mark range of post hoc statistical significance for nonsocial stimuli. For all panels: \*\*\**P* < 0.001, Mean ± SEM (shaded area/error bars) is presented. M, male; F, female; B, banana; P, peanut; H, hexanal; CA, clean air.

To quantify these differences and explore the detailed structure of the population code, we discretized neural responses of randomly selected groups of 10 cells from each mouse into 20 ms bins, and fitted maximum entropy models to the distributions of population activity patterns evoked by each of the presented stimuli (for each group of cells, we fitted both a first- and a second-order model, and used the one that gave higher cross-validation values, see Supplementary Fig. 3)^50,51^. We quantified the dissimilarity between the distributions of stimulus-evoked population responses (encoding distributions) using the Jensen-Shannon Divergence^52^,d(s_i_,s_j_)= D_js_[p(r|s_i_)||p(r|s_j_)], which measures in bits their distinguishability (*d* = 0 would indicate indistinguishable distributions, and *d* = 1 completely non-overlapping responses^53,54^; Fig. 3a bottom, see Online Methods). We calculated the dissimilarity between all pairs of encoding distributions, and averaged these distances across mice (presented in bits/s to give the rate of information about the stimulus identity; Fig. 3c). The block-diagonal structure of the dissimilarity matrix reflects a category-based organization of the population codebook in the mPFC: encoding distributions of the social cues were significantly more similar to each other than to any of the nonsocial odors, regardless of odor identity or valence (Fig. 3d). We further explored the divergence of stimulus encoding over time and found that while population activity patterns were indistinguishable during the baseline period, representations of social and nonsocial signals diverged within 2s following stimulus onset, and slowly returned to baseline levels after stimulus offset (Fig. 3e).

### Altered representation of social stimuli in the mPFC of Cntnap2^−/−^ mouse model of autism

Having observed distinct representations of social cues in the mPFC population code, we asked whether the mPFC representation of social signals is disrupted in animals that display impaired social function, such as those observed in autism spectrum disorder. In humans, mutations in the CNTNAP2/CASPR2 gene are strongly associated with ASD risk^55,56^, and patients that carry risk-associated variants of this gene show altered prefrontal connectivity^27^. Mice lacking the Cntnap2 gene present all of the core behavioral phenotypes of autism, as well as several associated neuronal phenotypes, including reduced cortical interneuron density^45^. We thus presented Cntnap2^−/−^ mice with the same set of social and nonsocial stimuli, using their age-matched wild-type littermates as controls (Cntnap2^+/+^; referred to as “wt” hereafter; *n*_Cntnap2−/−_=6, *n*_wt_=5). To characterize how population representations in the mPFC might undergo experience-dependent refinement, we compared neuronal responses to the stimuli in two consecutive recording sessions less than one week apart, for each mouse. In contrast with the previous experiment described in Fig. 1–3, in which mice were exposed to the odors prior to recording sessions, mice in the current experiment were habituated to the chamber but were not previously presented with the odor cues.

We recorded a total of 269 units in Cntnap2^−/−^ mice (133 on day 1 and 136 on day 2) and 237 in wt littermates (125 on day 1 and 112 on day 2, Supplementary Fig. 4). We found that while wt mice maintained a preference in unit response profile to social cues, this bias was lost in Cntnap2−/− mice (Fig. 4a,b). The distribution of unit selectivity shifted significantly between the two recording sessions in wt mice: the number of mixed-response units decreased between experimental sessions, whereas the percentage of units specifically responding to social or nonsocial stimuli, and the percentage of cue-specific units, increased (Fig. 4a and Supplementary Fig. 5). Cntnap2^−/−^ mice showed a similar decrease in the percentage of mixed-response units, but this was accompanied by an increase in the percentage of non-responsive units (from 44% on day 1 to 58% on day 2; Fig. 4a). The normalized magnitude of neuronal responses to social cues was significantly larger than for all nonsocial odors in wt mice, whereas this difference was significantly attenuated in Cntnap2^−/−^ mice (Fig. 4b). Importantly, the magnitude of stimulus-evoked neuronal responses was not correlated with behavioral locomotion in any of the genotypes or stimuli (Supplementary Table 1). In addition, the behavioral responses of Cntnap2^−/−^ mice to odor stimuli were indistinguishable from those of wt mice (Supplementary Fig. 6), consistent with previous work that demonstrated intact olfactory function in these mice^45^.

**Figure 4.**
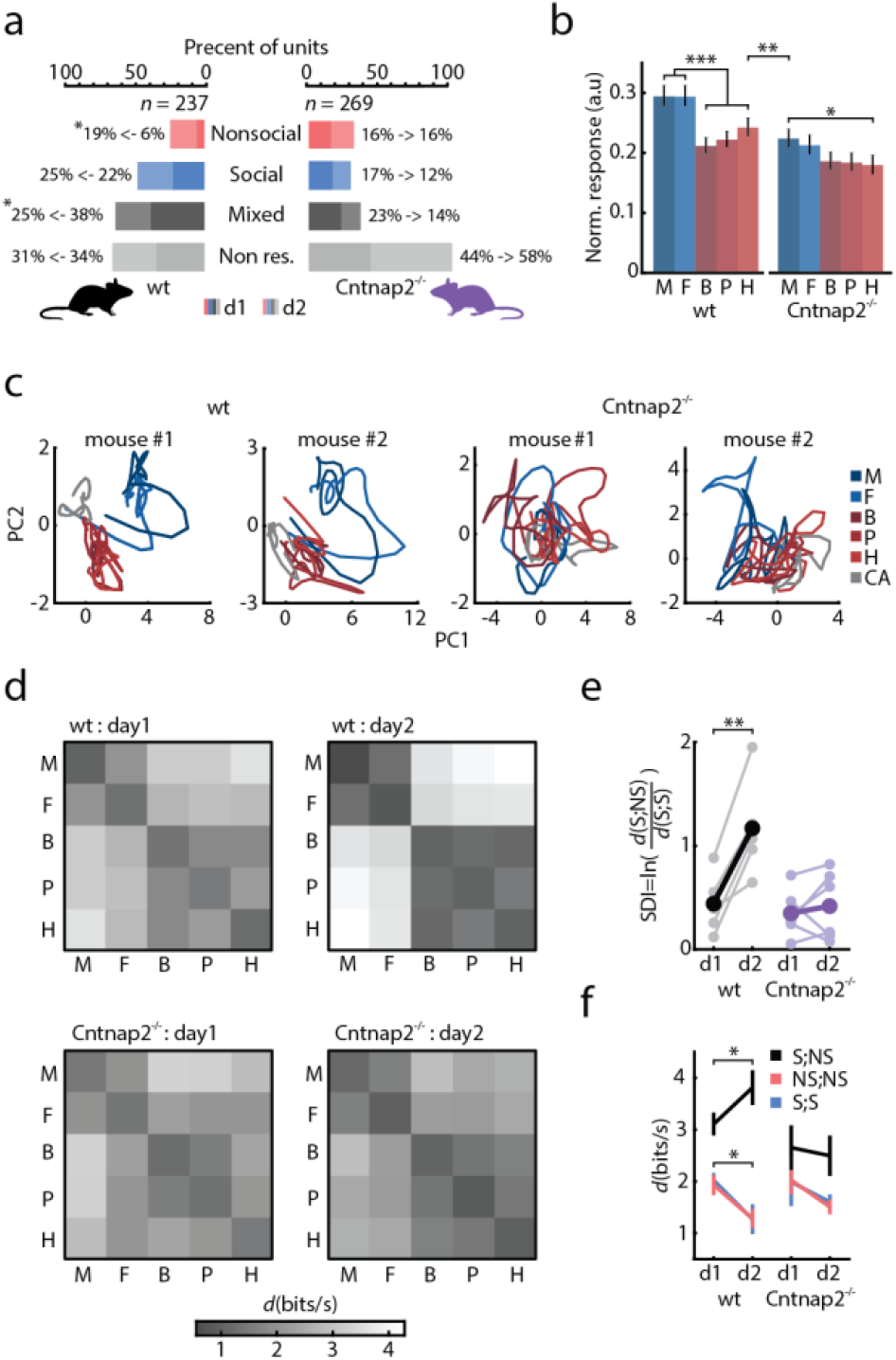
Altered dynamics of social representation in the Cntnap2^−/−^ mouse model of autism. **(a)** Distribution of response selectivity among recorded wt (*left*) and Cntnap2^−/−^ mPFC units (*right*) in two consecutive recording sessions. Dark colors indicate distribution on day 1 of recording, light colors mark distribution on day 2. Change in percentage of units between days is marked on the figure. *χ*^2^wt(3) = 11.957, *P* < 0.01; *χ*^2^_Cntnap2−/−(3)_ = 6.789, *P* = 0.079. Standardized residual analysis was used to determine post hoc significant changes in response categories between days (|standardized residual| > 2). **(b)** Overall unit tuning, presented as normalized response firing rate, for wt (*left*) and Cntnap2^−/−^ (*right*) mice. Mixed-design RM ANOVA with Bonferroni corrected post hoc comparisons. *F*genotype(1,288) = 10.653, *P* < 0.01; *F*stimulus(4,1152) = 14.097, *P* < 0.001; *F*genotype*stimulus(4,1152) = 2.291, *P* = 0.058. **(c)** Representative 2D projection of population activity trajectories during stimulus presentation for two wt (*left*) and two Cntnap2^−/−^ (*right*) mice during the second day of recording (see Online Methods). Colors represent odor identity. Here, the first two principal components accounted for 74% – 82% of the variance **(d)** Similarity matrices depicting the distance between population responses to stimuli in wt (*top*, *n* = 5) and Cntnap2^−/−^ (*bottom*; *n* = 6) mice, during the first (*left*) and second (*right*) recording days. **(e)** Distance-based social distinction index (SDI, see text) for wt and Cntnap2^−/−^ mice for two consecutive recording sessions. Higher index values indicate greater divergence between social and nonsocial stimuli. Bold lines depict mean values over mice, thin lines represent individual mice. Mixed-design RM ANOVA with Bonferroni corrections. *F*_genotype*day(1,9)_ = 14.05, *P* < 0.01; *F*_day(1,9)_ = 20.586, *P* < 0.01; *F*_genotype(1,9)_ = 5.14, *P* < 0.05. **(f)** Average dissimilarity between responses to odors (*d*) within categories (NS;NS and S;S) and between them (S;NS) for wt and Cntnap2^−/−^ mice, over two consecutive recording sessions. 2-way RM ANOVA with Bonferroni corrected post hoc comparisons. *F*dissimilarity*day (wt)(2,8) = 12.952, *P* < 0.01; *F*_dissimilarity (wt)(2,8)_ = 34.723, *P* < 0.001; *F*_day (wt)(1,4)_ = 2.925, *P* = 0.162; *F*dissimilarity*day (Cntnap2−/−)(2,10) = 0.5, *P* = 0.621; *F*_dissimilarity (Cntnap2−/−)(2,10)_ = 8.854, *P* < 0.01; *F*_day(Cntnap2−/−)(1,5)_ = 1.051, *P* = 0.352 (main effect of dissimilarity refers to differences between S;NS, NS;NS and S;S). For all panels: \**P* < 0.05, \*\**P* < 0.01, \*\*\**P* < 0.001. Mean ± SEM is presented. M, male; F, female; B, banana; P, peanut; H, hexanal; CA, clean air; Non.res, non-responsive; S, social; NS, nonsocial; d1, day 1; d2, day 2; SDI, social distinction index;

To examine the population activity in wt and Cntnap2^−/−^ mice, we again projected the population activity firing rates onto the first two principle components (as in Fig. 3b). Differences between wt and Cntnap2^−/−^ mice in population responses to the odors were clearly apparent in this low dimensional embedding: while wt trajectories prominently diverged in PC space based on social category (similar to data acquired in the original cohort shown in Fig. 3), Cntnap2^−/−^ trajectories showed no clear category-level separation (Fig. 4c).

### Social representations undergo experience-dependent refinement in wild-type but not in Cntnap2^−/−^ mice

In human ASD patients, disruption of plasticity-related processes has been proposed as an endophenotype of the disorder^57–59^. Consistent with these findings, several animal models of ASD display impairments in long-term synaptic plasticity^60,61^. We therefore measured the experience-dependent changes in mPFC population representations of odor stimuli by performing two recording sessions in each mouse, separated by 2–5 days. Again, we fitted a maximum entropy model to the population activity of randomly selected groups of 10 cells in each mouse, to each of the stimuli, for each of the two days, and compared the stimulus-evoked encoding distributions. While mPFC encoding in wt mice showed distinct social and nonsocial separation already in the first recording session, the distance between responses to odors from the two categories grew significantly larger on the second session (Fig. 4d, top), demonstrating a significant effect of experience. In contrast, the separation between the representations of social and nonsocial cues in Cntnap2^−/−^ mice was both less pronounced on the first session compared with wt, and did not improve on the following one (Fig. 4d bottom).

To quantify the separation between representation of social (S) and nonsocial (NS) stimuli for each mouse, we calculated a social distinction index, SDI=ln 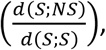, where *d*(S;NS) denotes the average distance between social and nonsocial stimuli and *d*(S;S) is the distance between social stimuli. The SDI values consistently and significantly increased between the first and second recording days for all wt mice. In contrast, no consistent change occurred in the Cntnap2^−/−^ group, and mean SDI values in these mice remained unchanged between sessions (Fig. 4e). We further found that in wt mice, the encoding distances within each category (*d*(S;S) and *d*(NS;NS)) decreased between recording sessions, while the inter-category distance *d*(S;NS) increased. In Cntnap2^−/−^ mice, however, no significant change occurred in either distance metric (Fig. 4f).

### Decoding of stimulus identity and category from mPFC population activity

To evaluate mPFC encoding of odor identity at the level of single trials, we used a maximum likelihood classifier based on the encoding models of the stimulus-evoked population activity. Models were trained on seven out of eight presentations of each stimulus and used to estimate the likelihood of odor identity for each one of the held-out test trials (all possible combinations of seven train trials and one test trial were calculated for each stimulus, see Online Methods). In wt mice, we could reliably decode both social and nonsocial odors from single-trial population activity (Fig. 5a,b, *left*). Averaged likelihood values in wt were similar for cues of the same 11 category, but lower for odors of the other category. Conversely, likelihood values in Cntnap2^−/−^ mice were similar for all odors regardless of social category (Fig. 5a,b).

**Figure 5.**
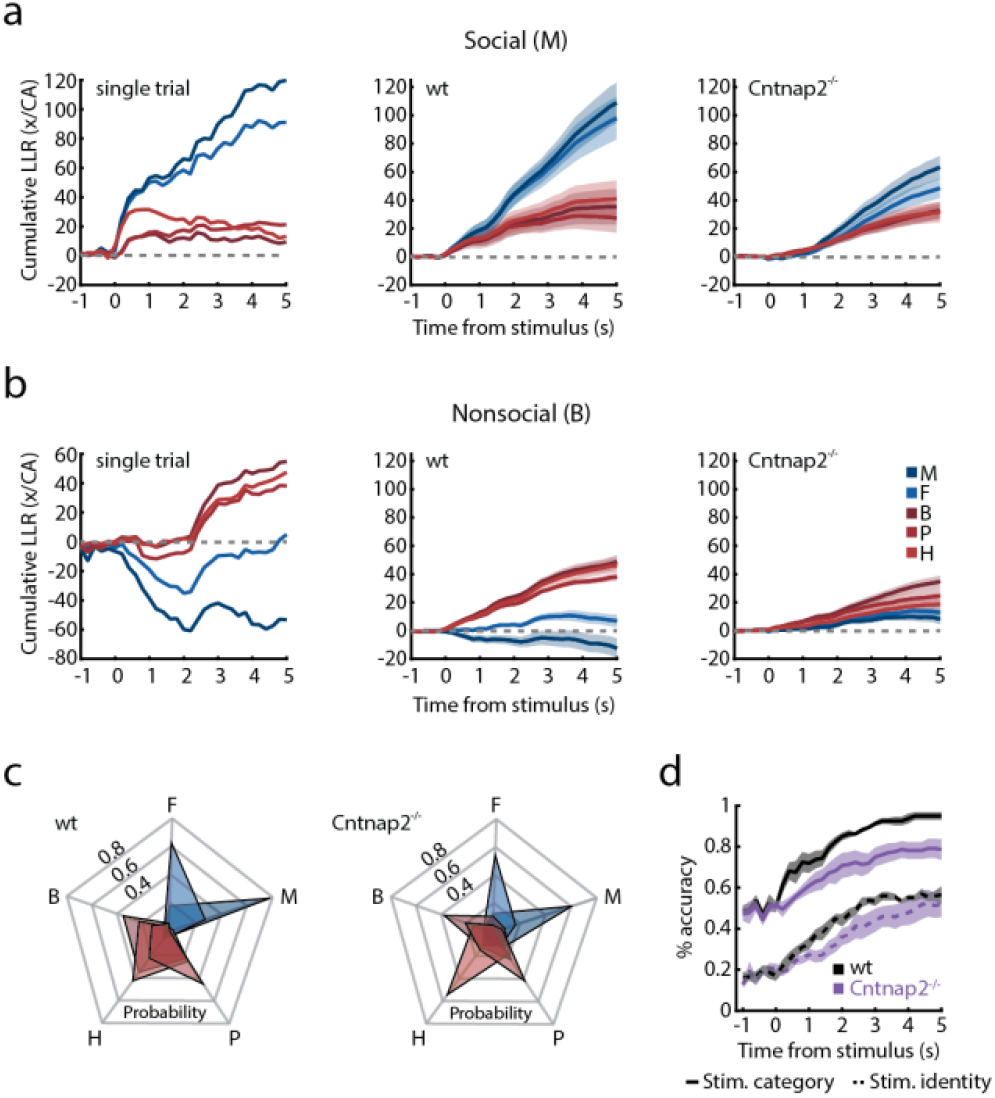
Decoding of stimulus identity and social category from mPFC population code. **(a)** *Left panel:* an example of the time-dependent cumulative log-likelihood ratio (LLR) of population responses to each stimulus and to clean air, on a single trial in which male odor was presented to a wt mouse. *Middle panel*: the average of the same LLR across all such trials over all wt mice. *Right panel*: the average across all such trials over all Cntnap2^−/−^ mice. **(b)** Time-dependent cumulative log-likelihood ratio (LLR) of each stimulus and clean air in trials where banana odor was presented (*left*: single trial data; *middle*: average for wt mice; *right*: average for Cntnap2^−/−^ mice). **(c)** Decoding performance for individual stimuli in wt (*left*) and Cntnap2^−/−^ (*right*) mice. Performance was summarized as the probability of classifying the presented stimulus as either one of all possible stimuli across all mice and trials. Colors and corresponding capital letters mark correct odor classification. **(d)** Cumulative accuracy of odor-based (*dashed lines*) and category-based (*solid lines*) decoders for wt (*black*) and Cntnap2^−/−^ (*purple*) mice. For all panels: Mean ± SEM are presented. M, male; F, female; B, banana; P, peanut; H, hexanal; Stim, stimulus.

We evaluated the performance of the decoders by the probability that the model of each stimulus would have the maximal likelihood value, given the presentation of a specific odor (Fig. 5c). Performance was well above chance for the correct odor in both wt and Cntnap2^−/−^ mice. However, while rare in wt mice, decoding errors between categories were common in Cntnap2^−/−^mice. The decoding results in wt mice demonstrated increased error rate within the nonsocial category, suggesting generalization of the representation of nonsocial odors (represented by the overlapping areas in Fig. 5c). To directly quantify the difference between genotypes, we next trained a stimulus-category decoder for social versus nonsocial odors and compared the results to those of the stimulus-identity decoder. Remarkably, decoding performance for individual odors was similar in wt and Cntnap2^−/−^ mice, whereas performance for odor category (S or NS) was significantly inferior in Cntnap2^−/−^ mice (Fig. 5d). These findings suggest that Cntnap2^−/−^ mice have specific deficits in odor categorization, rather than in encoding of stimulus identity.

### Increased neuronal noise in mPFC activity of Cntnap2^−/−^ mice correlates with altered social representation

Findings from human ASD patients^22,62^ and mouse models of autism^35–37^ implicate altered cortical E/I ratio during brain development in the pathophysiology of the disorder^32^–34,63. This is in line with earlier reports of reduced cortical interneuron numbers in Cntnap2^−/−^ mice^45,64^. Consistently, we found that the mean baseline firing rates of mPFC units recorded in Cntnap2^−/−^ mice were significantly higher than those recorded in their wt littermates (Fig. 6a). Furthermore, the pairwise correlations between units in Cntnap2^−/−^ mice were lower than in wt mice. During stimulus presentation, pairwise correlations showed a greater increase in wt mice compared with their Cntnap2^−/−^ littermates (Supplementary Fig. 7a). We quantified the correlation at the level of the entire population by the second-order connected information^65^, and found that correlations in wt mice increased during stimulus presentation while they remained unchanged in Cntnap2^−/−^ mice (Supplementary Fig. 7b).

Since E/I imbalance was hypothesized to result in elevated cortical “noise”^66^, we calculated the average variability of population firing patterns in baseline (ongoing) neuronal activity (Online Methods). We found that this measure of noise was significantly higher in Cntnap2^−/−^ mice compared to their wt littermates (Fig. 6b-c; note that with the exception of one mouse, the noise measure of the two genotypes were completely non-overlapping). An example of 2D projections of population activity traces during ongoing activity is presented in Fig. 6b, overlaid on the population traces during stimulus presentation. Multiple regression analysis revealed that the elevated baseline noise levels in Cntnap2^−/−^ mice could not be linearly predicted by either elevated baseline unit firing rates, nor by behavioral locomotion levels (Supplementary Fig. 7c,d). Remarkably, we found that across genotypes, baseline neuronal noise showed a strong negative correlation with SDI values (Fig. 6d). In contrast, SDI values did not correlate with average unit firing rates in either genotype (Supplementary Fig. 7e). In line with our results above, no significant correlation between noise level and SDI values was found in the first recording day, but a strong correlation emerged in the second recording session, when SDI values increased in wt mice but remained low in Cntnap2^−/−^ mice (Fig. 6d). To further explore this finding, we calculated the change in SDI values between recording sessions for each mouse (ΔSDI). Strikingly, ΔSDI values in individual mice were strongly correlated with baseline cortical noise, such that elevated noise levels were predictive of decreased experience-dependent refinement of mPFC social representation (Fig. 6e).

**Figure 6.**
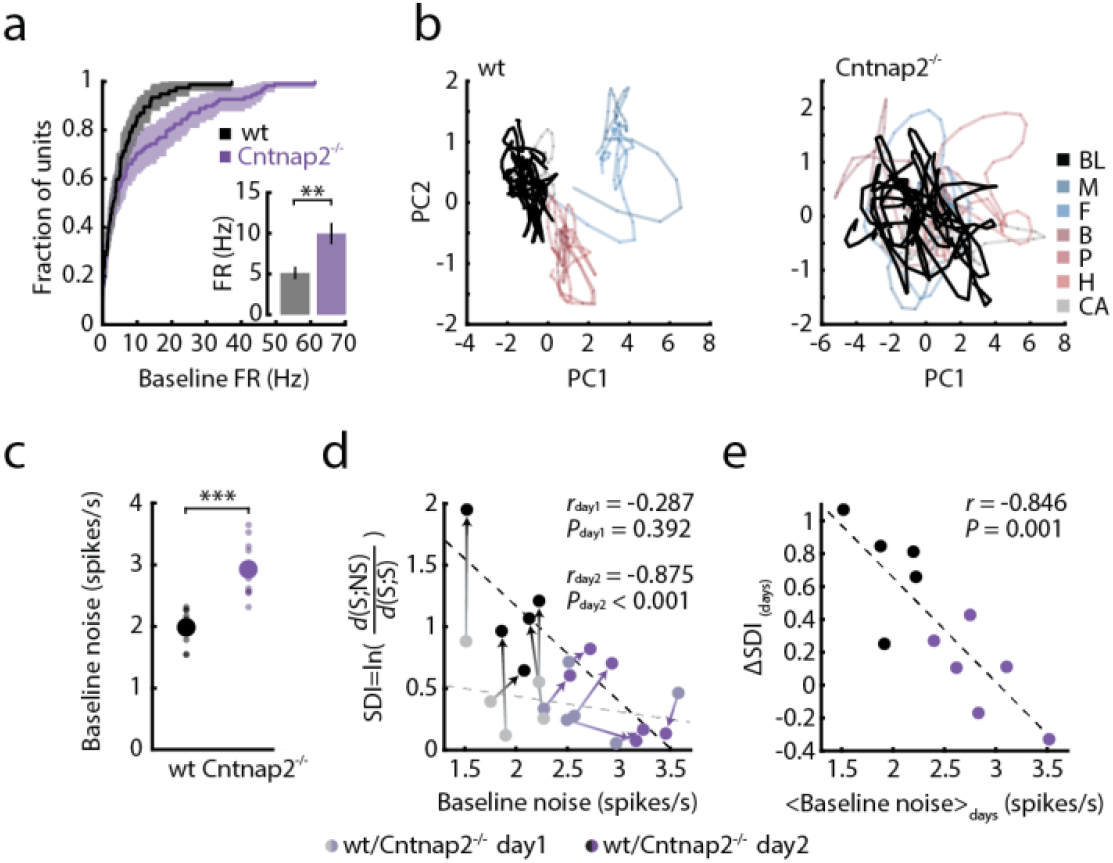
Elevated neural noise correlates with deficits in social processing in Cntnap2^−/−^ mice. **(a)** Cumulative distribution of baseline firing rates of units recorded in wt (*black*) and Cntnap2^−/−^ (*purple*) mice. Shaded area marks 95% confidence interval. Inset depicts average firing rate values. Student’s t-test, *t*_(164)_ = 3.182, *P* < 0.01. **(b)** Representative 2D projection of neural trajectories of baseline activity (*black line*) in wt (*left*) and Cntnap2^−/−^ (*right*) mice. Traces are overlaid on corresponding trajectories of stimulus-evoked activity in the same mice (light color lines; see Fig. 4c). **(c)** Average baseline noise as calculated by the integrated movement in population activity space during pre-stimulus baseline (see Online Methods and panel b for 2D projection of this measurement). Student’s t-test, *t*_(20)_ = 5.902, *P* < 0.001 **(d)** Correlation between baseline noise level and SDI values for wt (*black*) and Cntnap2^−/−^ mice (*purple*), for the first (*bright colors*) and second (*dark colors*) recording sessions. Correlation was calculated across genotypes for each recording day. Corresponding correlation values are indicated. Circles represent individual mice; arrows connect values of the same mouse from first to second recording day. (**e**) Correlation between average baseline noise level and change in SDI values between recording days for wt and Cntnap2^−/−^ mice. Correlation was calculated across genotypes and corresponding values are marked on figure. For all panels: Mean ± SEM is presented (note: some SEM in panels c-e are smaller than marker size). \*\**P* < 0.01, \*\*\**P* < 0.001. M, male; F, female; B, banana; P, peanut; H, hexanal; CA, clean air; SDI, social distinction index.

### Discussion

In this study, we investigated the neural encoding of social chemosensory signals in the prefrontal cortex of freely behaving mice. We found robust tuning to social stimuli in mPFC unit activity and distinct population responses to social versus nonsocial signals. Repeated exposure of mice to the same set of cues revealed experience-dependent refinement of these representations between days. These results provide the first evidence for representation of defined social chemosensory cues in the mouse mPFC. Consistent with previous reports of processing of salient sensory signals in the mPFC^17,18,67,68^, we found that stimulus category (social/nonsocial), rather than individual odor identity, is predominantly represented in the mPFC neural code. Recent studies exploring stimulus encoding in olfactory cortical regions reported no difference in single-cell or population responses to social versus nonsocial odors^12,69^. The social categorization we observe in the mPFC signifies an additional tier of processing, possibly relying on converging information from odor-driven activity in several long-range synaptic inputs to the mPFC (e.g., piriform cortex^70^, orbitofrontal cortex^71,72^), as well as on inputs from brain regions encoding social context and incentive salience, such as the amygdala^3,48^, VTA^73^ and ventral hippocampus^74,75^.

Examining the responses to sensory cues across two separate recording sessions, we found that population activity patterns underwent significant refinement in wt mice. These findings expand upon recent work describing experience-dependent divergence of conspecific sex representations in the hypothalamus^76^ and medial amygdala^77^. In contrast to these regions, in which representations of sex specific signals diverge with experience, population activity in the mPFC seems to categorize cues based on their social or nonsocial properties, and this contrast is further refined with experience whereas within-category representations grow similar with time.

Our findings further show that mPFC activity in the Cntnap2^−/−^ mouse model of autism displays reduced selectivity to social stimuli and loss of social categorization, while retaining information about the identity of individual odors. Most strikingly, Cntnap2^−/−^ mice lacked the robust experience-dependent changes observed in the population code of wt mice. Impairments in short and long-term plasticity processes were previously described in both human ASD patients^57–59^ and animal models of the disorder^60,61^, but were not explored on the circuit level, nor linked to the neuronal representation of social information. The loss of experience-based refinement of odor category representation in Cntnap2^−/−^ mice might be linked with the role of CNTNAP2 protein in targeting AMPA and NMDA receptors to post-synaptic membranes^78,79^. Furthermore, recent work by Lazaro et al. (“Reduced prefrontal synaptic connectivity and disturbed oscillatory population dynamics in the CNTNAP2 model of autism”, *bioRxiv 2018*) demonstrates reduced dendritic spine and synapse densities in the mPFC of Cntnap2^−/−^ mice, providing a potential mechanism for the functional impairments we describe. The loss of these circuit-level plasticity processes in Cntnap2−/− mice might contribute to reduced selectivity in the mPFC representation of salient social cues and constitute a potential neuronal mechanism for the social impairment displayed by these mice^45^.

Which network-wide changes might underlie these deficits? Leading theories of ASD attribute its symptoms to increased cortical excitation/inhibition ratio during brain development^32^. Our findings of elevated firing rates and altered correlation structure in Cntnap2^−/−^ mice are consistent with this theory and with previous reports of decreased density of cortical inhibitory interneurons in this model^45,80^. In line with these findings, a recent study demonstrated that elevation of inhibitory neuron activity in the mPFC of Cntnap2^−/−^ mice leads to restoration of social behavioral responses^81^, suggesting that the behavioral deficits in Cntnap2^−/−^ mice may be intimately linked with the altered mPFC representations we observe here.

Importantly, we also find elevated population activity noise in Cntnap2^−/−^mice, similar to findings in human ASD patients^31,82,83^. The strong negative correlation we observe between baseline noise and social category representations suggests that noise plays an important role in the failure of mPFC activity in Cntnap2^−/−^ mice to appropriately represent social stimuli and drive synaptic plasticity processes. These deficits might lead to an impaired ability to adaptively respond to relevant cues during social interactions. Taken together, our results present new insights into the encoding of social information in the mPFC, and provide a neurophysiological perspective on the association between altered neocortical processing and social dysfunction in autism.

## Acknowledgements

This work was supported by grants from the Simons Foundation, the European Research Council (ERC-2013-StG OptoNeuromod 337637, and ERC 311238 NEURO-POPCODE), the Israel Science Foundation, the Israel-US Binational Science Foundation, the Adelis Foundation, the Lord Sieff of Brimpton Memorial Fund and the Candice Appleton Family Trust. O.Y. is supported by the Gertrude and Philip Nollman Career Development Chair. E.S. is the incumbent of the Joseph and Bessie Feinberg Professorial Chair.

## Author contributions

D.R.L and O.Y. designed the study, D.R.L and A.W. built the experimental setup, D.R.L performed all experiments and analyzed unit activity and behavioral data. D.R.L, T.T., OY, and ES performed population coding analyses. M.K. contributed to behavioral data analysis. D.R.L., O.Y., T.T, and E.S. wrote the manuscript. D.R.L and T.T. contributed equally to this manuscript.

## Online Methods

### Animals

Animals used for this study were 3–5 months old male C57BL/6J (*n* = 6; Envigo, Rehovot, Israel), Cntnap2^−/−^ (*n* = 6) and Cntnap2^+/+^ (*n* = 5) mice (courtesy of Prof. Elior Peles of the Weizmann Institute of Science). The Cntnap2 knockout mice were previously back-crossed to a C57BL/6J background for at least 10 generations^45^, and maintained by heterozygote breeding. Mice were kept on a 12-h light/dark cycle with food and water *ad libitum*, and tested during the dark phase. All procedures described in this paper were approved by the Weizmann Institute Animal Care and Use Committee (IACUC).

### Stereotaxic surgery and microwire array implantation

Mice were anesthetized with an intraperitoneal injection of Ketamine–Xylazine mixture (80 mg/kg Ketamine, 10 mg/kg Xylazine), placed into a stereotaxic frame (David Kopf Instruments) and kept under 1.5% isoflurane anesthesia throughout the procedure. Microwire electrode arrays were implanted in the infralimbic region of the medial prefrontal cortex (distance from Bregma according to the Allen brain atlas: AP: +1.97; ML ±0.3 counter balanced between mice; DV = −3.0), and secured to the skull using Metabond (Parkell) and dental acrylic. Analgesic (Buprenorphine, 0.05 mg/kg) was provided immediately post-surgery. Mice were placed in an individual cage and allowed 2-weeks to recover before initiation of experimental trials. Locations of implanted drives were validated in all experimental animals using an electrolytic lesion (see histology section below and Supplementary Fig. 1).

### In-vivo electrophysiological recordings

Multi-electrode drive consisted of a graded electrode bundle of 16 microwires (25-μm diameter straightened tungsten wires; Wiretronic Inc.), attached to an 18-pin dual row connector (Mill-Max, Oyster Bay, NY). Unit signals were amplified using a HS-18-CNR-LED unity-gain headstage amplifier, filtered (600–6,000 Hz), digitized at 32 kHz and stored using the Digital Lynx hardware and Cheetah software acquisition system (Neuralynx Inc.).

### Odor infusion apparatus

The apparatus consists of a transparent polycarbonate chamber (10cm × 15 cm × 15cm), connected to a custom-made 7-odor olfactometer plugged into a 1/8” odor inlet in the chamber floor. Odor stimuli were placed in individual polytetrafluoroethylene (PTFE) vials, each directed to the chamber through a separate tube system converging onto a designated PTFE hub at the inlet odor port. One-way check valves were placed in each odor path to prevent back-flow of odors.

Odors were infused via constant airflow stream directed through alternating solenoids controlled by a MOSFET Electronic driver. Odor alternation occurred within ∼12ms (as measured using a pressure sensor, see Supplementary Fig. 1a). Air from the chamber was constantly cleared using a vacuum system in order to maintain constant pressure and clear odor residue throughout the experiment. In/out airflows were controlled using four 24VDC pressure pumps (Conlog Ltd. Israel) and fine-tuned using a built-in valve. The kinetics of odor concentration in the chamber were assessed using a VOC meter (MiniRAE Lite; RAE systems, San Jose CA. see Supplementary Fig. 1b). Odor concentrations showed a sharp increase immediately after stimulus onset, continued to increase throughout stimulus infusion, and slowly decreased back to baseline levels (a decrease of an order of magnitude in concentration was measured within ∼60 seconds from stimulus off). All air pumps were isolated inside a sound attenuating box designed to minimize noise levels. All pipes, inlets and odor tubes were either constructed of or coated with PTFE to prevent odor contamination.

The setup was back-lit with a planar infra-red (IR) LED array (880nm, 1Vision Ltd., Israel), allowing high-contrast recording and analysis of mouse behavior. The IR backlight was isolated from the behavioral chamber with a transparent conductive mask (Holland Shielding Systems B.V., the Netherlands) to minimize electrical noise in recorded channels. Two buffered 1.3MP monochromatic infrared triggered CMOS cameras (Mightex Systems), as well as the Neuralynx IR camera were used to record the experiment from a top and side view simultaneously, allowing for analysis of behavior with high-temporal resolution alongside the electrophysiological data. All components of the setup were controlled using a National Instruments data logger (NI USB-6353, National Instruments, Austin, TX), and a custom-written Matlab program. All events in the odor delivery setup were logged on the Neuralynx system using digital TTL inputs.

### Odor stimuli and experimental procedure

Social cues consisted of soiled bedding and 50µl of urine collected and pooled from 10 male (M) or 10 female (F) adult C57BL/6J mice, in order to minimize the effect of individual cues between experimental repetitions. Nonsocial odor stimuli were: natural Banana (B) and Peanut (P) extracts (Sensale, Ramat-Gan, Israel) and a monomolecular Hexanal (H) odorant (Sigma-Aldrich), all diluted 10^−2^ in DDW on the morning of each experiment.

At the beginning of each experimental day, mice were connected to the electrophysiological tether and placed in the chamber for 15–30 minutes to allow habituation to the setup and stabilization of electrophysiological signals. The experimental procedure initiated with three minutes of baseline recording followed by odor presentation trials. Each trial consisted of 10 seconds of clean air, followed by 5 seconds of odors infusion and an additional 60 seconds of clean air infusion to clear the chamber of odor residue (see Fig 1a). Each experiment consisted of 40 such trials (8 trials for each of the 5 selected stimuli: M/F/B/P/H). Trials were pseudo-randomized to prevent multiple consecutive presentations of a single odor, and interleaved with eight additional clean air trials using the same trial design (to account for possible changes in airflow and sound due to solenoid switching). After the end of odor delivery trials, ongoing spontaneous activity was recorded for at least 5 additional minutes. Vacuum pump was constantly activated throughout the experiment (including habituation and baseline times), and clean air was constantly infused into the chamber with the exception of odor delivery times. All experiments were done under a dim ambient light of 3 Lux.

For C57BL/6J experiments, mice were repeatedly exposed to the odor stimuli prior to initiation of experiments. Mice were used for two recording session separated more than a month apart. For experiments involving Cntnap2^−/−^ and Cntnap2^+/+^ mice, mice were never before exposed to odor stimuli prior to the first experimental day. In this experiment, recording session were conducted 2 or 5 days apart, with inter-session gaps similarly distributed between the two groups.

## Data analysis

### Behavioral analysis

Recorded videos were automatically analyzed frame by frame, using custom-written MATLAB scripts (version 2017a, MathWorks, Natick, MA). Videos of experimental sessions were segmented using a fixed-threshold, and body contour was distinguished from electrophysiological head stage and tether using erosion and dilation procedures. The center of mass (CoM) of the mouse was then determined for future analysis. Locomotion values were calculated by integrating the Euclidean distances (absolute values) between pairs of CoM values in consecutive frames, over a period of 5 s during odor presentation or immediately beforehand (for baseline measurements). Behavioral attention response and orientation to odor infusion were scored manually frame-by-frame. Behavioral data was averaged per mouse (across trials and sessions) unless otherwise indicated. One mouse was excluded from behavioral analysis on a single recording session due to technical issue with the recorded video file. All analyses were conducted by a trained observer, blind to stimulus identity and mouse genotype.

### Analysis of electrophysiological unit data

Neuronal data was sorted using Plexon OfflineSorter 3.2.4 (Plexon, Dallas, TX, USA), based on principal component analysis of spike waveform and inter-spike interval (ISI). Prior to sorting, the raw signal from all simultaneously recorded channels was averaged and subtracted from each channel using a custom Matlab script, in order to remove global electrical noise artifacts. To determine unit responsivity, evoked firing rates were calculated for the 5 seconds of stimulus presentation and compared to baseline firing rates during the preceding 5 seconds of baseline recordings. Response *Z*-score was calculated across repetitions per stimulus, per unit, and |Z-score| ≥ 2 threshold was used to determine responsive units and response specificity. A range of additional thresholds were also tested to provide further validation for the consistency of our results (See Supplementary Fig. 2e). Minimum empirical standard deviation (calculated across the entire data set for each stimulus) was used for Z-score analysis of units that were silent during baseline recordings, but responded during stimulus presentation (< 2 % of instances). Normalized response magnitude used for the unit-tuning analysis was evaluated as: 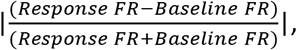, averaged per stimulus, across repetitions, and used in absolute values unless indicated otherwise. Firing rates and Z-scored PSTHs were calculated in 250 ms bins, averaged across repetitions per unit, and then averaged across all units responding to each specific stimulus (only units significantly increasing their firing rate in response to stimulus presentations were used for PSTH analysis). Single unit firing rates, used for baseline FR comparisons between genotypes, were collected and averaged across 60 s of baseline recordings conducted before the initiation of experimental procedure. Due to habituation in unit responses, the first 5 presentations of each stimulus (out of 8 trials) were used for the single unit analysis.

### Population activity analysis

To study population coding at a fine temporal resolution, we discretized population activity patterns into 20 ms bins, where the activity of the units at time bin *t* was given by a binary vector 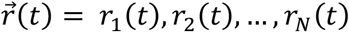, where *r*_*i*_ = 1(0) denotes whether neuron *i* spiked in that bin. Since estimating the encoding distribution 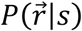 directly from the data is impractical due to under-sampling (see Supplementary Fig. 3), we constructed for each time window a model 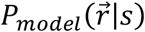 of the distribution of neural responses as a function of time, based on the minimal models that have the correct firing rates of individual units, 〈*r*_*i*_(*t*)〉, and pairwise correlations between them 〈*r_i_*(*t*)*r_j_*(*t*)〉 (where 〈 〉, denote average over stimulus presentations) known as stimulus dependent maximum entropy models^51^. For each population recorded from each animal and for each stimulus, we fitted two models: (1) the maximum entropy model based only on the time dependent firing rates, giving the conditionally independent population model, 
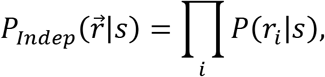
which assumes no correlations between units, and (2) a stimulus-dependent second-order maximum entropy (ME2) model that also takes into account the time dependent correlations between units, as previously described^50,54^. The ME2 model is known to take the form:
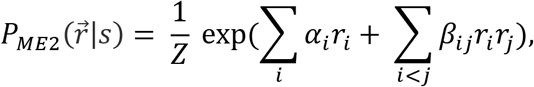
where {α_*i*_, *β_ij_*} are Lagrange multipliers that were fitted so that the averages {〈*r_i_*(*t*)〉, 〈*r_i_*(*t*)*r_j_*(*t*)〉} of the model agree with experimental data, and *Z* is a normalization term or the partition function. We then estimated the likelihood of held out test data for each of the models, in order to choose the model which provided the best fit to the data (see Supplementary Fig. 3a). The chosen model was then used in the analysis of the neural population activity patterns. In all cases, the models gave a highly accurate description of the data, which was superior to those based on the empirically sampled responses (see Supplementary Fig 3c). Despite the response habituation observed over repeated cue presentation for single unit responses, using subsampling of trials for construction population models (either early trials 1–4, or later trials 5–8) did not affect the population analysis results.

To quantify the dissimilarity of stimulus-evoked population activity patterns, the “distance” between two stimuli was quantified as the dissimilarity between their encoding distributions, calculated by the Jensen-Shannon divergence^49^: 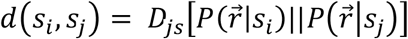. The Jensen-Shannon divergence is a symmetrized version of the Kullbak-Leibler divergence, which measures in bits how distinguishable two distributions are, yielding 0 for identical distributions and 1 for non-overlapping distributions^52^.

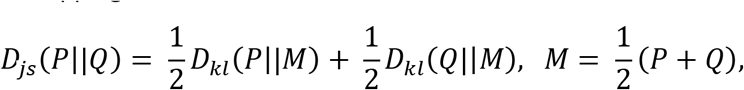

For each animal, the encoding distributions models were fitted to 30 randomly selected groups of ten units. In order to evaluate self-distance (*d*(x,x)) we fitted two models of the encoding distribution of each stimulus, using half of the trials (odd/even trials) as training data for each model, and then calculated the D_js_ between them: 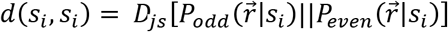. A single mouse was removed from this analysis (presented in Fig. 3c-e) due to insufficient number of simultaneously recorded units (7 units).

For analysis of population neural trajectories, spike trains were discretized in non-overlapping bins of 150 ms and convolved with a Gaussian kernel (width: 150 ms). Trial-averaged population activity vectors representing the instantaneous state of the system 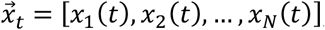, (N is the number of units), were then projected onto the first two principal components using PCA.

To evaluate cortical noise levels, we calculated the average fluctuations of population activity vectors during ongoing (baseline) activity before stimulus presentations: 
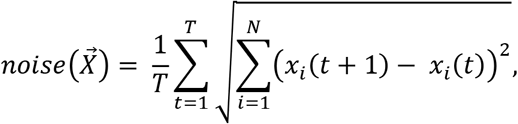
where T is the length of ongoing segment and 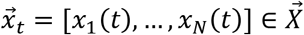 is the population activity vector at time *t*. To avoid bias due to difference in population size, we used groups of ten randomly selected units (20 groups in each mouse).

### Single-trial decoding of stimulus identity/category

To decode stimulus identity and stimulus category, we constructed maximum likelihood classifiers using encoding models of the entire population of simultaneously recorded units. Models were trained on seven randomly selected trials out of eight experimental trials for each stimulus and tested on the 8^th^ trial. As described earlier, the best model for each training set was used to estimate the likelihood of observing each one of the stimuli, given the population responses in the held-out test trials. For each trial, we calculated the cumulative log likelihood ratio (LLR) of each of the stimulus encoding models and the model and the clean air response over time:

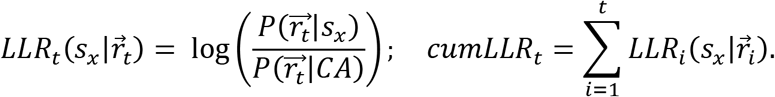

The trial was then classified according to the model that gave the highest likelihood at the end of stimulus presentation, 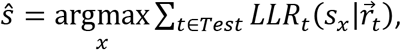 and decoder performance was defined as the probability of choosing each one of the possible stimuli given the presentation of a specific odor. The category-based decoder was trained and tested on a combination of trials of different stimuli from the same category (social/nonsocial). Data were chosen such that the train and the test sets of the two categories would consist of the same number of trials.

#### Statistical analysis

All analyses and subsequent statistical tests were performed using Matlab (The Mathworks Inc.) or Statistica software (StatSoft Inc.). Bonferroni corrections or Dunnett’s test were used when appropriate to correct for post hoc comparisons. Levene’s test was used to assess equality of variances, and statistical parameters were adjusted accordingly when needed. All statistical tests presented in this manuscript are two-tailed. Details of specific statistical designs and appropriate tests are described for each analysis in the appropriate figure legend. Statistical significance of post hoc analysis is marked on the relevant figure panels.

#### Data availability

The data that support the findings of this study and custom written analysis codes are available from the corresponding author upon reasonable request.

#### Histology

Mice were deeply anesthetized using an intraperitoneal injection of Ketamine–Xylazine mixture (160 mg/kg Ketamine, 20 mg/kg Xylazine) and the locations of implanted electrodes were marked with electrolytic lesions (unipolar 100 µA current for 5 s, for each polarity). Twenty minutes following the lesion procedure, mice were further anesthetized using Pentobarbital (130 mg/kg^−1^, i.p.), and then transcardially perfused with ice-cold phosphate buffered saline (PBS, pH 7.4) followed by 4% paraformaldehyde solution. Brains were extracted, post-fixed overnight at 4 °C in 4% PFA, and then moved to 30% sucrose solution for at least 48 hours. Coronal sections (35μm) were acquired using a microtome (Leica Microsystems) and collected in a cryoprotectant solution (25% glycerol, 30% ethylene glycol in PBS, pH 6.7). Sections were stained with a nucleic acid dye to better visualize lesion location (DAPI, 1:10,000), mounted on gelatin-coated slides, dehydrated and embedded in DABCO mounting medium (Sigma). Tiled overview images (X10) were acquired using a LSM 700 confocal microscope (Zeiss), and electrode locations were recorded.

---

**Supplementary Figure 1.**
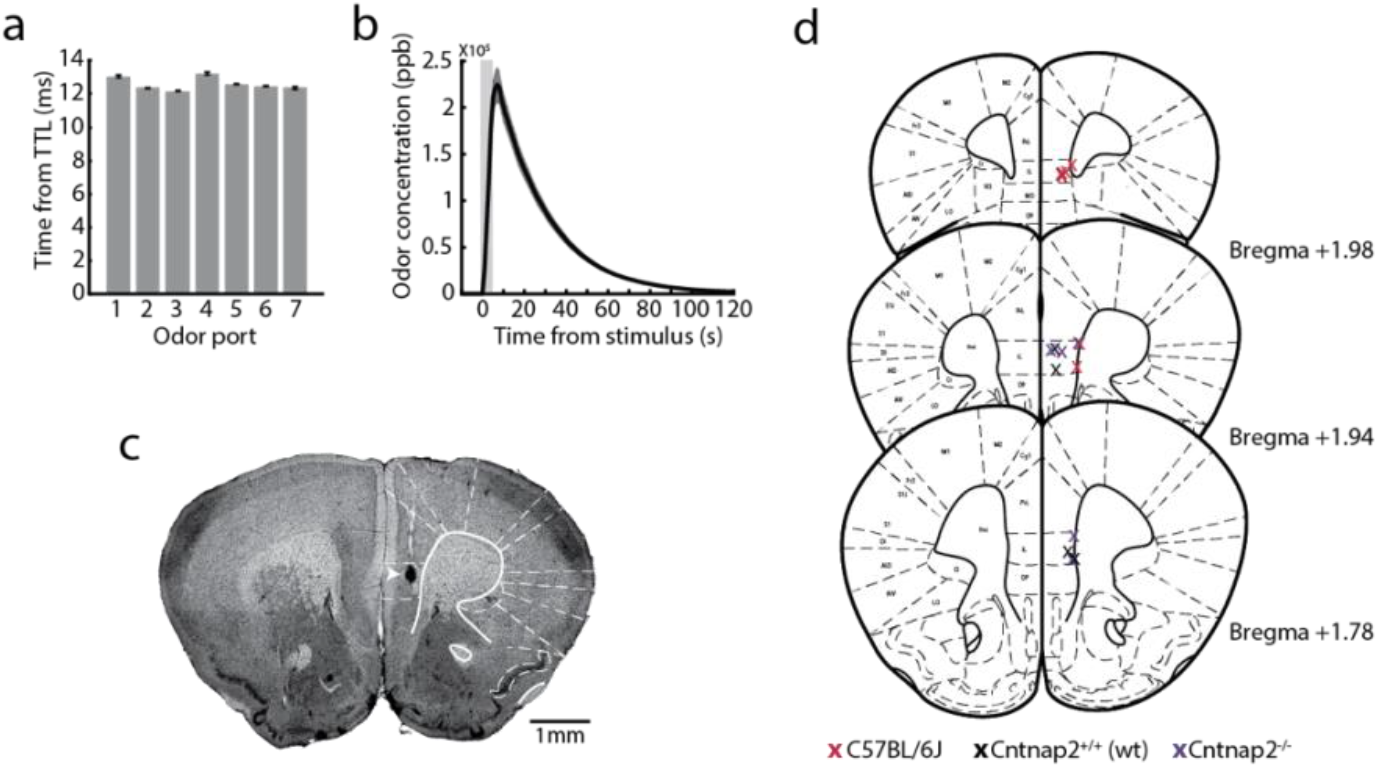
Olfactometer calibrations and microarray electrode location. **(a)** Latency to odor infusion from each of seven available odor ports. A pressure sensor was used to determine initiation of airflow into the chamber following TTL input indicating the opening of the appropriate odor solenoid. Each solenoid was tested 5 consecutive times. Mean ± SEM is presented. **(b)** Change in odor concentration at the center of the chamber. Measurements were taken using a volatile organic compound (VOC) meter following 5 sec infusion of vapor from a 70% ethanol solution followed by infusion of clean air (5 repetitions). Shaded area marks stimulus presentation times. Mean ± SEM is presented. **(c)** Representative image depicting the location of an electrolytic lesion used to verify electrode position in the mPFC. Arrow indicates lesion location in the infalimbic cortex. **(d)** Schematic representation of electrode placement in recorded mice from all experimental groups. Location was determined using the most ventral end of localization lesion or electrode track.

**Supplementary Figure 2.**
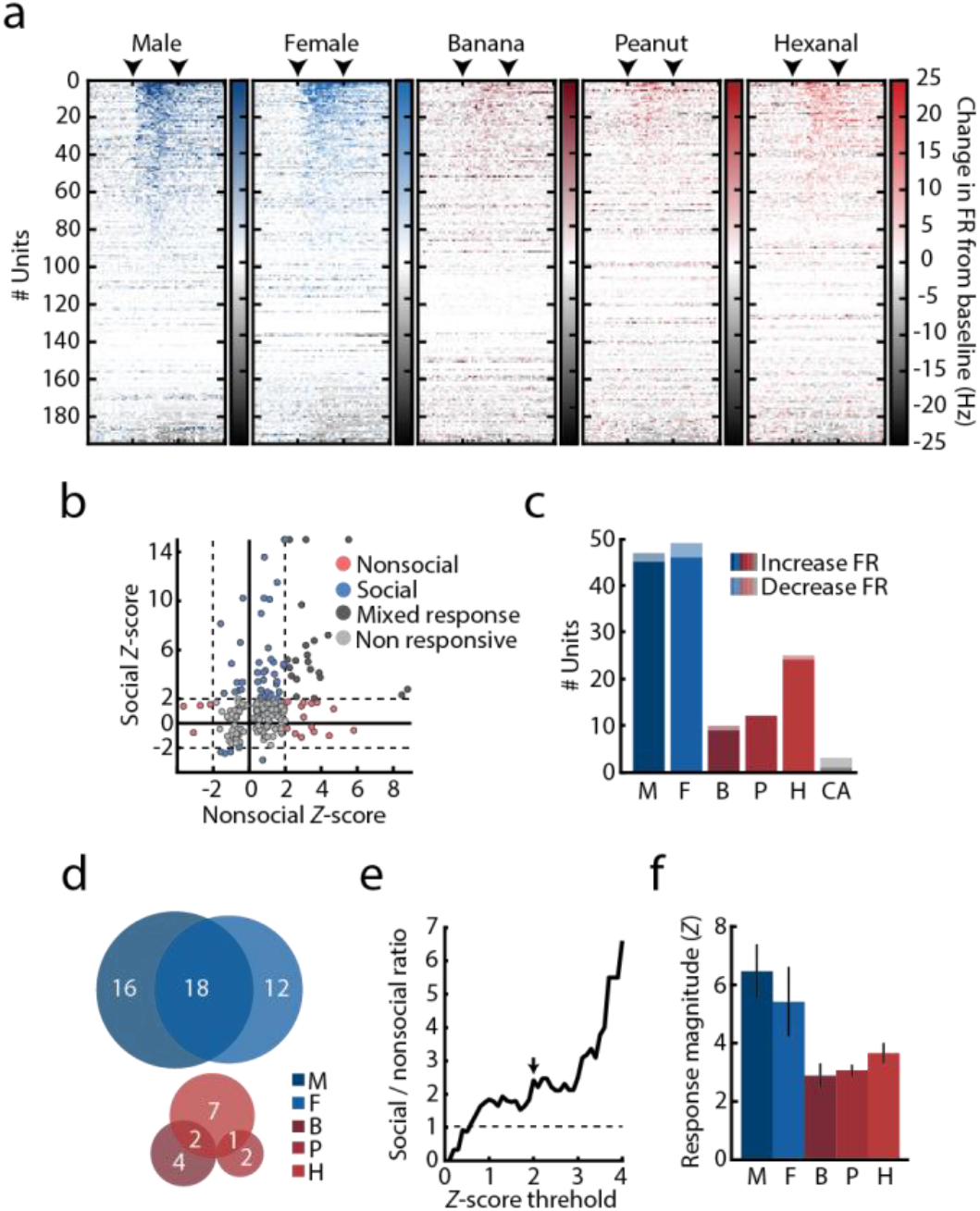
Social tuning in the mPFC unit responses. **(a)** Stimulus-evoked response across all recorded units per stimulus, sorted by response magnitude as calculated by the absolute change in firing rate from baseline. Color gradient represents the normalized change in firing rate calculated over 250ms bins. Arrowheads mark the time of stimulus onset and offset. **(b)** Response *Z*-score distribution calculated for all recorded units in response to social (M/F) and nonsocial (B/P/H) cues. Circles represent maximum response of individual units to each stimulus category. Color code represents response specificity. Z score threshold (*Z* = 2) is represented by a dashed line. Units with *Z* > 15 were assigned with *Z* = 15 for presentation purposes. **(c)** Number of units significantly increasing (*dark*) and decreasing (*bright*) their firing rates in response to each presented stimulus. Color represents stimulus identity. **(d)** Stimulus specificity overlap within social (*top*) and nonsocial (*bottom*) units. Number of units in each category is indicated on the figure. Circle sizes are scaled to the number of units responding to each stimulus. **(e)** Relative ratio between social and nonsocial units, calculated using a continuous range of *Z* score thresholds. Arrow represents *Z* = 2. Note that the number of social units consistently exceeds that of nonsocial units starting at *Z* > 0.6 **(f)** Average response magnitude for all presented stimuli for units significantly increasing their firing rate in response to stimulus presentations. Mean ± SEM is presented. One way ANOVA, *F*_stimulus(4,131)_ = 1.603, *P* = 0.177. For all panels: M, male; F, female; B, banana; P, peanut; H, hexanal; CA, clean air.

**Supplementary Figure 3.**
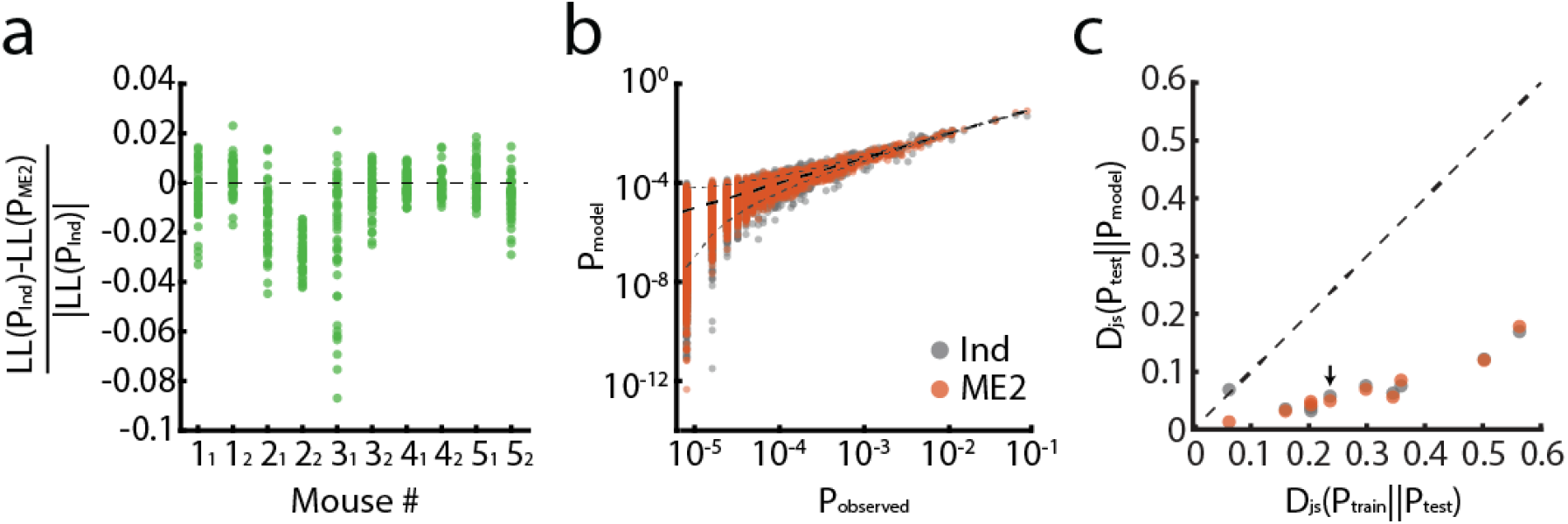
Maximum entropy models accurately describe the encoding distributions of the stimuli. **(a)** Normalized difference between the log-likelihood values of the pairwise maximum entropy model (ME2) and conditionally independent model, for each mouse. Models were trained over seven trials of a specific stimulus and tested on one held-out trial per stimulus. Each dot corresponds to one held-out trial for one specific stimulus (6 stimuli × 8 trials = 48 dots per mouse). Positive values indicate larger likelihood for the independent model over the ME2; the most likely model for each trial was then used for decoding analysis (see Fig. 5). **(b)** The empirical probabilities of population activity patterns of cells recorded in one mouse in response to one odor are plotted against the probabilities predicted by different models (gray dots, independent model; orange dots, ME2 model). Each dot corresponds to a single activity pattern observed during the experiment. The funnel marked by the dashed grey line indicates 99% confidence interval of the empirical measurement. Black dashed line shows equality. **(c)** The Jensen–Shannon divergences between the empirical joint probability distribution of activity patterns and the different models – ME2 (orange) and conditionally independent (gray). Black line indicates equality of the distance of the models from the test data, and the distance between the training and test data. Models were trained using randomly chosen 1750 samples, similar to the number of training data sample used for the decoding analysis (7 trials of 5 seconds each). Analysis was done using all recorded units from each mouse (up to 20 units) and the mean over ten randomly chosen training sets is plotted. While no model is consistently better than the other in capturing the distribution across all mice, both models clearly outperform the empirical model. Arrow indicates the example mouse shown in panel **b**.

**Supplementary Figure 4.**
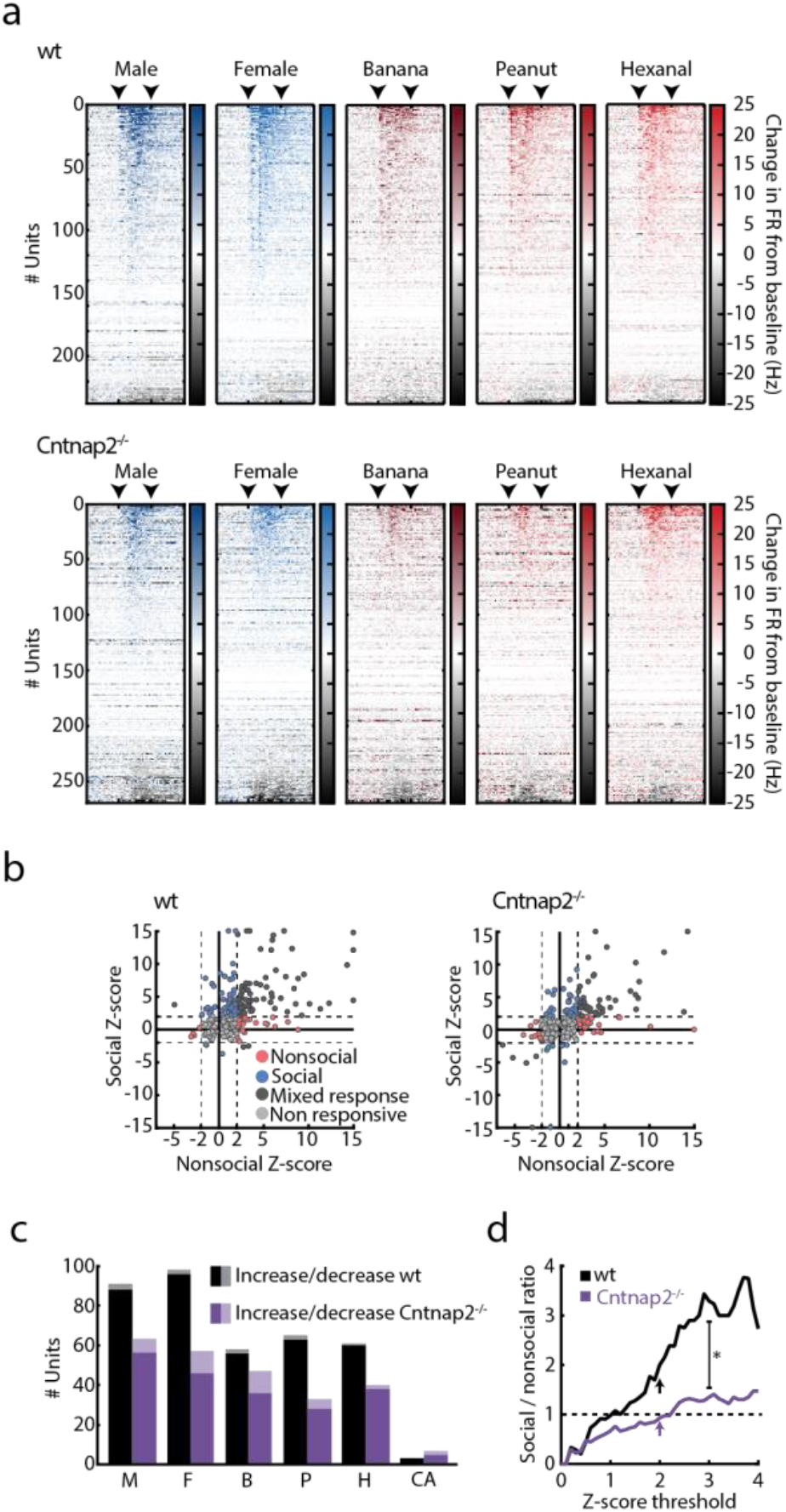
Altered response patterns to social and nonsocial stimuli in the mPFC of Cntnap2^−/−^ mice. **(a)** Stimulus-evoked responses across all recorded units per stimulus, sorted by response magnitude as calculated by the change in firing rate from baseline for wt (*top*) and Cntnap2^−/−^ (*bottom*) mice. Color gradient represents the change in firing rate from baseline, calculated over 250ms bins. Arrows mark the time of stimulus onset and offset. **(b)** Response Z-score distribution calculated for all recorded units in wt (*left*) and Cntnap2^−/−^ mice (*right*) in response to social (M/F) and nonsocial (B/P/H) cues. Circles represent maximum response of individual units to each stimulus category. Color code represents response specificity. *Z* score threshold (|*Z*| = 2) is represented by a dashed line. Units with *Z* > 15 were assigned with *Z* = 15 for presentation purposes. **(c)** Number of units significantly increasing (*dark*) and decreasing (*bright*) their firing rates in response to each presented stimulus in wt (*black*) and Cntnap2^−/−^ (*purple*) mice. **(d)** Relative ratio between social and nonsocial units, calculated using a continuous range of *Z* score thresholds for wt (*black*) and Cntnap2^−/−^ (*purple*) mice. Arrows represent *Z* = 2. Linear regression analysis (0≤ *Z* ≤3), *F*_wt(1,30)_ = 1088.42, *P*< 0.001, *F*_Cntnap2−/−(1,30)_ = 652.294, *P* < 0.001, *B*_wt_ = 1.106, *B*_Cntnap2−/−_ = 0.425, with non-overlapping 95% confidence intervals. For all panels: \**P* < 0.05, M, male; F, female; B, banana; P, peanut; H, hexanal; CA, clean air.

**Supplementary Figure 5.**
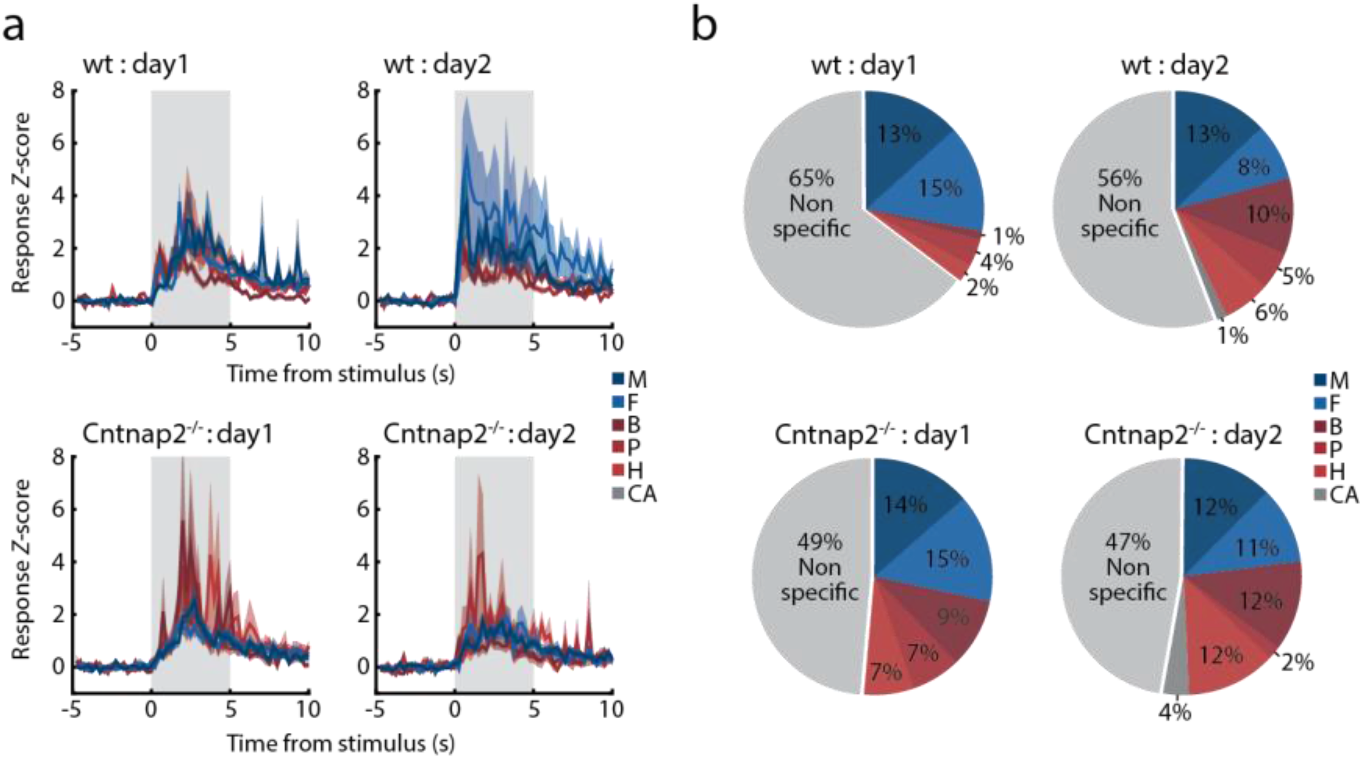
Experience-dependent changes in stimulus-evoked unit responses. **(a)** Stimulus-evoked PSTHs portraying mean increase in response *Z*-score of cue-responsive units in the first (*left*) and second (*right*) recording sessions, for wt (*top*) and Cntnap2^−/−^ (*bottom*) mice. Color code represent stimulus identity. Shaded areas mark stimulus presentation time. Mean ± SEM is presented. **(b)** Stimulus specificity among cue responsive units in the first (*left*) and second (*right*) recording sessions, in wt (*top*) and Cntnap2−/− (*bottom*) mice. Colors represent stimulus identity. For all panels: M, male; F, female; B, banana; P, peanut; H, hexanal; CA, clean air.

**Supplementary Figure 6.**
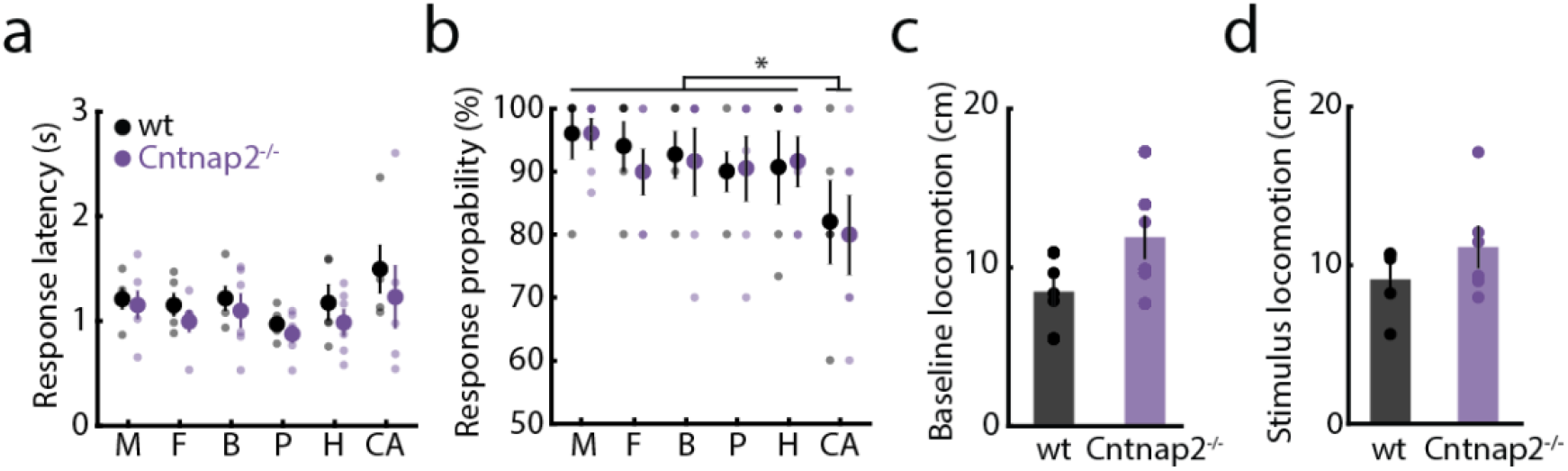
Behavioral responses to odor presentation in wt and Cntnap2^−/−^ mice. **(a)** Average latency to odor-evoked orientation responses for wt (*black*) and Cntnap2^−/−^ (*purple*) mice. Circles represent individual mice. Mixed-design RM ANOVA. *F*_genotype(1,9)_ = 0.959, *P* = 0.352; *F*_stimulus(5,45)_ = 2.449, *P* <0.05; *F*_genotype*stimulus(5,45)_ = 0.163, *P* = 0.974 **(b)** Mean probability of odor-evoked orientation responses for wt (*black*) and Cntnap2^−/−^ (*purple*) mice, across all odors. Circles represent individual mice. Mixed-design RM ANOVA. *F*_genotype(1,9)_ = 0.040, *P* = 0.844; *F*_stimulus(5,45)_ = 3.304, *P* < 0.05 with Dunnett’s test against clean air; *F*_genotype*stimulus(5,45)_ = 0.115, *P* = 0.988 **(c)** Baseline behavioral locomotion levels of wt and Cntnap2^−/−^ mice. Mann-Whitney U Test, *U* = 7, *P* = 0.171 **(d)** Mean behavioral locomotion during odor presentation, averaged across all presented odors. Mann-Whitney U Test, *U* = 10, *P* = 0.411. For all panels: Mean ± SEM is presented. *n*_wt_ = 5, *n*_cntnap2−/−_ = 6, \**P* < 0.05. M, male; F, female; B, banana; P, peanut; H, hexanal; CA, clean air.

**Supplementary Figure 7.**
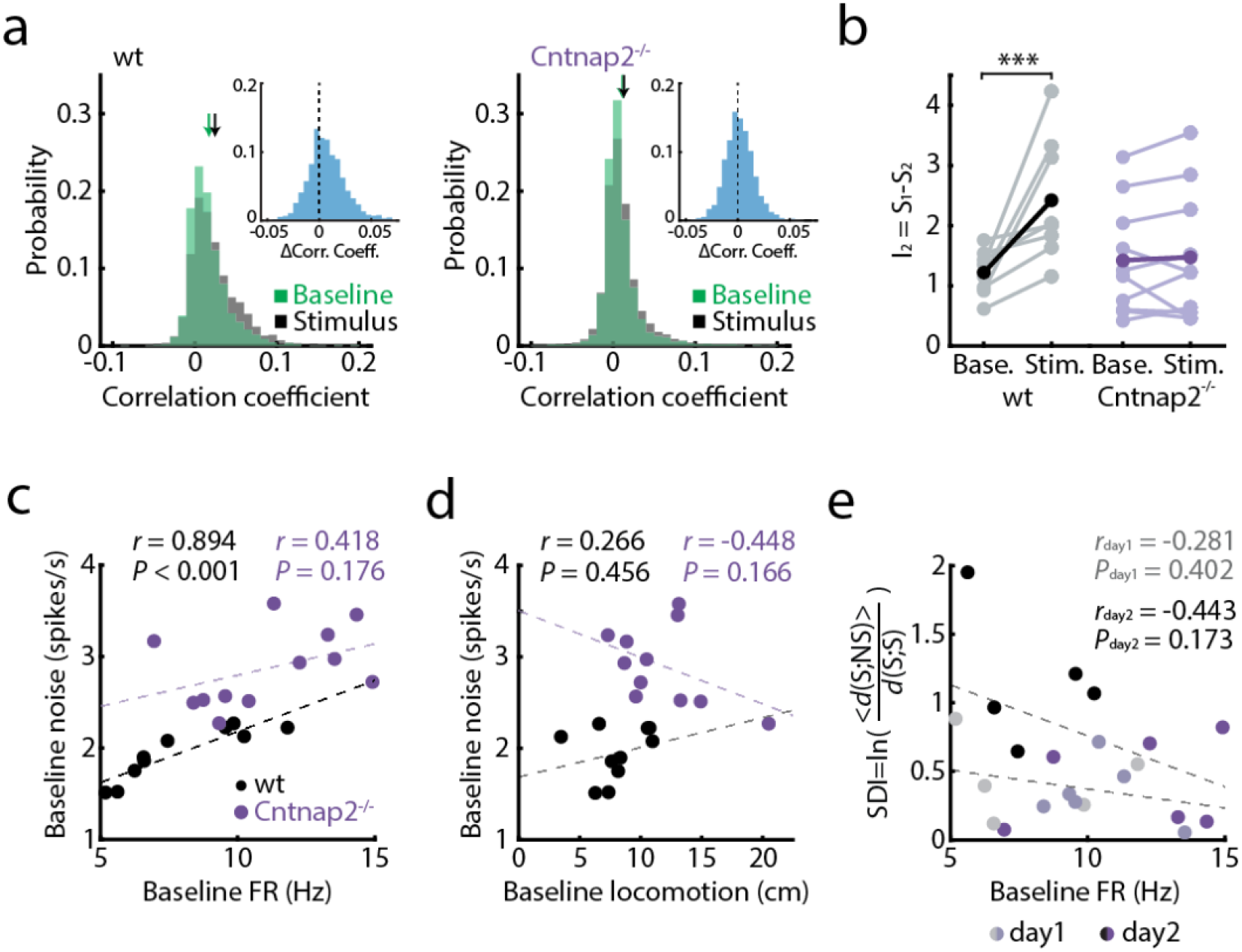
Elevated baseline neural noise in Cntnap2^−/−^ mice is not predicted by behavioral locomotion or baseline firing rate. **(a)** Distribution of the Pearson correlations between all neuron pairs, estimated using a 5 s window during baseline (green) and stimulus presentation (grey) in units recorded from the mPFC of wt (*left*) and Cntnap2^−/−^ mice (*right*). Arrows mark the average correlation value in each experimental phase. Insets depict the distribution of differences in pairwise correlations between stimulus and baseline periods. **(b)** Information in pairwise interactions (second order connected information) during baseline and stimulus periods for wt (*black*) and Cntnap2^−/−^ (*purple*) mice. Higher values indicate greater contribution of pairwise correlations in the neuronal code. Bold lines depict mean values, lighter lines represent individual mice. Mixed-design RM ANOVA with post hoc Bonferroni correction. *F*_genotype*phase(1,17)_ = 12.746, *P* < 0.01; *F*_phase (1,17)_ = 15.114, *P* < 0.01; *F*_genotype(1,17)_ = 1.14, *P* = 0.30. One wt mouse and one Cntnap2−/− mouse which showed aberrant values (> 5 standard deviations from the mean) were removed from this analysis. **(c-e)** Correlation between baseline FR and baseline noise levels **(c)**, between baseline behavioral locomotion and baseline noise levels **(d)**, and between baseline FR and SDI values in recording day 1 (light) and recording day 2 (dark) **(e)** for wt (*black*) and Cntnap2^−/−^ (p*urple*) mice. Correlation in **(c,d)** is calculated for values collected from both recording days. Corresponding values and statistical significance are marked on figure. Multiple linear regression with FR and locomotion as linear predictors of noise was performed: *F*_wt(2,7)_ = 16.480, *P* < 0.01, with FR as significant predictor *P* < 0.001. *F*_Cntnap2−/−(2,9)_ = 1.680, *P* = 0.240. Circles represent individual mice in a single recording session. \*\*\**P* < 0.01; For all panels: Base., baseline; Stim., stimulus; d1, day1; d2, day2.

**Supplementary Table 1.** Unit stimulus-evoked response magnitude is not correlated with stimulus-evoked change in behavioral locomotion. Response *Z*-score values were calculated for all units responding to each odor in each mouse in each recording session (in absolute values, averaged across trials). Pearson’s correlations were then used to assess correlation of unit response magnitude with the corresponding change in behavioral locomotion during infusion of odor stimuli compared to baseline.

|  | Male | Female | Banana | Peanut | Hexanal |
| --- | --- | --- | --- | --- | --- |
| **Wild-type** | $r = 0.299$<br>$P = 0.401$ | $r = 0.187$<br>$P = 0.605$ | $r = -0.146$<br>$P = 0.688$ | $r = 0.097$<br>$P = 0.789$ | $r = 0.246$<br>$P = 0.494$ |
| **Cntnap2<sup>-/-</sup>** | $r = -0.027$<br>$P = 0.937$ | $r = -0.019$<br>$P = 0.956$ | $r = -0.317$<br>$P = 0.342$ | $r = -0.087$<br>$P = 0.799$ | $r = 0.462$<br>$P = 0.153$ |

## References

1. Chen, P. & Hong, W. Neural Circuit Mechanisms of Social Behavior. Neuron 98, 16–30 (2018).

2. Lin, D. Y., Zhang, S.-Z., Block, E. & Katz, L. C. Encoding social signals in the mouse main olfactory bulb. Nature 434, 470–477 (2005).

3. Bergan, J. F., Ben-Shaul, Y. & Dulac, C. Sex-specific processing of social cues in the medial amygdala. Elife 3, e02743 (2014).

4. Anderson, D. J. Circuit modules linking internal states and social behaviour in flies and mice. Nat. Rev. Neurosci. 17, 692–704 (2016).

5. Franklin, T. B. et al. Prefrontal cortical control of a brainstem social behavior circuit. Nat. Neurosci. 20, 260–270 (2017).

6. Hong, W., Kim, D.-W. & Anderson, D. J. Antagonistic Control of Social versus Repetitive Self-Grooming Behaviors by Separable Amygdala Neuronal Subsets. Cell 158, (2014).

7. Lin, D. et al. Functional identification of an aggression locus in the mouse hypothalamus. Nature 470, 221–226 (2011).

8. Euston, D. R., Gruber, A. J. & McNaughton, B. L. The Role of Medial Prefrontal Cortex in Memory and Decision Making. Neuron 76, 1057–1070 (2012).

9. Adolphs, R. The social brain: neural basis of social knowledge. Annu Rev Psychol 60, 693–716 (2009).

10. Bicks, L. K., Koike, H., Akbarian, S. & Morishita, H. Prefrontal cortex and social cognition in mouse and man. Front. Psychol. 6, 1–15 (2015).

11. Watson, K. K. & Platt, M. L. Social signals in primate orbitofrontal cortex. Curr Biol 22, 2268–2273 (2012).

12. Kim, Y. et al. Mapping social behavior-induced brain activation at cellular resolution in the mouse. Cell Rep. 10, 292–305 (2015).

13. Zhou, T. et al. History of winning remodels thalamo-PFC circuit to reinforce social dominance. Science 357, 162–168 (2017).

14. Murugan, M. et al. Combined Social and Spatial Coding in a Descending Projection from the Prefrontal Cortex. Cell 171, 1663–1677.e16 (2017).

15. Lee, E. et al. Enhanced Neuronal Activity in the Medial Prefrontal Cortex during Social Approach Behavior. J. Neurosci. 36, 6926–6936 (2016).

16. Wang, F. et al. Bidirectional control of social hierarchy by synaptic efficacy in medial prefrontal cortex. Science (80-.). 334, 693–7 (2011).

17. Otis, J. M. et al. Prefrontal cortex output circuits guide reward seeking through divergent cue encoding. Nature 543, 103–107 (2017).

18. Milad, M. R. & Quirk, G. J. Neurons in medial prefrontal cortex signal memory for fear extinction. Nature 420, 70–74 (2002).

19. Rigotti, M. et al. The importance of mixed selectivity in complex cognitive tasks. Nature 497, 1–6 (2013).

20. Parthasarathy, A. et al. Mixed selectivity morphs population codes in prefrontal cortex. Nat. Neurosci. (2017). doi:10.1038/s41593-017-0003-2

21. Zikopoulos, B. & Barbas, H. Changes in prefrontal axons may disrupt the network in autism. J. Neurosci. 30, 14595–609 (2010).

22. Oblak, A., Gibbs, T. T. & Blatt, G. J. Decreased GABAA receptors and benzodiazepine binding sites in the anterior cingulate cortex in autism. Autism Res. 2, 205–19 (2009).

23. Lombardo, M. V. et al. Atypical neural self-representation in autism. Brain 133, 611–624 (2010).

24. Solso, S. et al. Diffusion tensor imaging provides evidence of possible axonal overconnectivity in frontal lobes in autism spectrum disorder toddlers. Biol. Psychiatry 79, 676–684 (2016).

25. Carper, R. A. & Courchesne, E. Localized enlargement of the frontal cortex in early autism. Biol. Psychiatry 57, 126–133 (2005).

26. Catani, M. et al. Frontal networks in adults with autism spectrum disorder. Brain 139, 616–630 (2016).

27. Scott-Van Zeeland, A. A. et al. Altered Functional Connectivity in Frontal Lobe Circuits Is Associated with Variation in the Autism Risk Gene CNTNAP2. Sci. Transl. Med. 2, 56ra80–56ra80 (2010).

28. Dinstein, I. et al. Disrupted Neural Synchronization in Toddlers with Autism. Neuron 70, 1218–1225 (2011).

29. Hahamy, A., Behrmann, M. & Malach, R. The idiosyncratic brain: distortion of spontaneous connectivity patterns in autism spectrum disorder. Nat. Neurosci. 18, 302–309 (2015).

30. Baum, S. H., Stevenson, R. A. & Wallace, M. T. Behavioral, perceptual, and neural 26 alterations in sensory and multisensory function in autism spectrum disorder. Progress in Neurobiology 134, (2015).

31. Dinstein, I. et al. Unreliable Evoked Responses in Autism. Neuron 75, 981–991 (2012).

32. Rubenstein, J. L. R. & Merzenich, M. M. Model of autism: increased ratio of excitation/inhibition in key neural systems. Genes. Brain. Behav. 2, 255–267 (2003).

33. Nelson, S. B. & Valakh, V. Excitatory/Inhibitory Balance and Circuit Homeostasis in Autism Spectrum Disorders. Neuron 87, 684–698 (2015).

34. Lee, E., Lee, J. & Kim, E. Excitation/Inhibition Imbalance in Animal Models of Autism Spectrum Disorders. Biological Psychiatry 81, 838–847 (2017).

35. Yizhar, O. et al. Neocortical excitation/inhibition balance in information processing and social dysfunction. Nature 477, 171–178 (2011).

36. Han, S. et al. Autistic-like behaviour in Scn1a+/-mice and rescue by enhanced GABA-mediated neurotransmission. Nature 489, 385–390 (2012).

37. Chao, H.-T. et al. Dysfunction in GABA signalling mediates autism-like stereotypies and Rett syndrome phenotypes. Nature 468, 263–269 (2010).

38. Tabuchi, K. et al. A neuroligin-3 mutation implicated in autism increases inhibitory synaptic transmission in mice. Science (80-.). 318, 71–76 (2007).

39. Peñagarikano, O. et al. Exogenous and evoked oxytocin restores social behavior in the Cntnap2 mouse model of autism. Sci. Transl. Med. 7, (2015).

40. Mei, Y. et al. Adult restoration of Shank3 expression rescues selective autistic-like phenotypes. Nature 530, (2016).

41. Peixoto, R. T. et al. Early hyperactivity and precocious maturation of corticostriatal circuits in Shank3B−/− mice. Nat. Neurosci. 19, (2016).

42. Wang, X. et al. Altered mGluR5-Homer scaffolds and corticostriatal connectivity in a Shank3 complete knockout model of autism. Nat. Commun. 7, 11459 (2016).

43. Nagode, D. A. et al. Abnormal Development of the Earliest Cortical Circuits in a Mouse Model of Autism Spectrum Disorder. Cell Rep. 18, 1100–1108 (2017).

44. Ey, E., Leblond, C. S. & Bourgeron, T. Behavioral profiles of mouse models for autism spectrum disorders. Autism Research 4, 5–16 (2011).

45. Peñagarikano, O. et al. Absence of CNTNAP2 leads to epilepsy, neuronal migration abnormalities, and core autism-related deficits. Cell 147, 235–246 (2011).

46. Stowers, L. & Kuo, T.-H. Mammalian pheromones: emerging properties and mechanisms 27 of detection. Curr Opin Neurobiol 34, 103–109 (2015).

47. Matsuo, T. et al. Genetic dissection of pheromone processing reveals main olfactory system-mediated social behaviors in mice. Proc. Natl. Acad. Sci. U. S. A. 112, E311–20 (2015).

48. Root, C. M., Denny, C. A., Hen, R. & Axel, R. The participation of cortical amygdala in innate, odour-driven behaviour. Nature 515, 269–273 (2014).

49. Kobayakawa, K. et al. Innate versus learned odour processing in the mouse olfactory bulb. Nature 450, 503–508 (2007).

50. Schneidman, E., Berry, M. J., Segev, R. & Bialek, W. Weak pairwise correlations imply strongly correlated network states in a neural population. Nature 440, 1007–1012 (2006).

51. Granot-Atedgi, E., Tkačik, G., Segev, R. & Schneidman, E. Stimulus-dependent Maximum Entropy Models of Neural Population Codes. PLoS Comput. Biol. 9, (2013).

52. Lin, J. Divergence Measures Based on the Shannon Entropy. IEEE Trans. Inf. Theory 37, 145–151 (1991).

53. Schneidman, E. Towards the design principles of neural population codes. Curr. Opin. Neurobiol. 37, 133–140 (2016).

54. Tkacik, G., Granot-Atedgi, E., Segev, R. & Schneidman, E. Retinal metric: A stimulus distance measure derived from population neural responses. Phys. Rev. Lett. 110, (2013).

55. Rodenas-Cuadrado, P. et al. Characterisation of CASPR2 deficiency disorder – a syndrome involving autism, epilepsy and language impairment. BMC Med. Genet. 17, 8 (2016).

56. Alarcon, M. et al. Linkage, association, and gene-expression analyses identify CNTNAP2 as an autism-susceptibility gene. Am J Hum Genet 82, (2008).

57. Bourgeron, T. From the genetic architecture to synaptic plasticity in autism spectrum disorder. Nat. Rev. Neurosci. 16, 551–563 (2015).

58. Swartz, J. R., Wiggins, J. L., Carrasco, M., Lord, C. & Monk, C. S. Amygdala habituation and prefrontal functional connectivity in youth with autism spectrum disorders. J. Am. Acad. Child Adolesc. Psychiatry 52, 84–93 (2013).

59. Guiraud, J. A. et al. Differential habituation to repeated sounds in infants at high risk for autism. Neuroreport 22, 1 (2011).

60. Auerbach, B. D., Osterweil, E. K. & Bear, M. F. Mutations causing syndromic autism define an axis of synaptic pathophysiology. Nature 480, 63–68 (2011).

61. Contractor, A., Klyachko, V. A. & Portera-Cailliau, C. Altered Neuronal and Circuit Excitability in Fragile X Syndrome. (2015). doi:10.1016/j.neuron.2015.06.017 62.

62. Bozzi, Y., Provenzano, G. & Casarosa, S. Neurobiological bases of autism-epilepsy comorbidity: A focus on excitation/inhibition imbalance. European Journal of Neuroscience (2017). doi:10.1111/ejn.13595 63.

63. Dickinson, A., Jones, M. & Milne, E. Measuring neural excitation and inhibition in autism: Different approaches, different findings and different interpretations. Brain Research 1648, 277–289 (2016).

64. Vogt, D. et al. Mouse Cntnap2 and Human CNTNAP2 ASD Alleles Cell Autonomously Regulate PV+ Cortical Interneurons. Cereb. Cortex 1–12 (2017). doi:10.1093/cercor/bhx248

65. Schneidman, E., Still, S., Berry, M. J. & Bialek, W. Network Information and Connected Correlations. Phys. Rev. Lett. 91, 238701 (2003).

66. Shadlen, M. N. & Newsome, W. T. The variable discharge of cortical neurons: implications for connectivity, computation, and information coding. J. Neurosci. 18, 3870–96 (1998).

67. Hardung, S. et al. A Functional Gradient in the Rodent Prefrontal Cortex Supports Behavioral Inhibition. Curr. Biol. 27, 549–555 (2017).

68. Alexander, W. H. & Brown, J. W. Medial prefrontal cortex as an action-outcome predictor. Nat Neurosci 14, 1338–1344 (2011).

69. Iurilli, G. & Datta, S. R. Population Coding in an Innately Relevant Olfactory Area. Neuron 93, 1180–1197.e7 (2017).

70. Vertes, R. P. Differential projections of the infralimbic and prelimbic cortex in the rat. Synapse 51, 32–58 (2004).

71. Price, J. L. et al. Olfactory input to the prefrontal cortex. Olfaction A Model Syst. Comput. Neurosci. 101–120 (1991).

72. Schoenbaum, G. & Eichenbaum, H. Information coding in the rodent prefrontal cortex. I. Single-neuron activity in orbitofrontal cortex compared with that in pyriform cortex. J. Neurophysiol. 74, 733–50 (1995).

73. Gunaydin, L. A. et al. Natural neural projection dynamics underlying social behavior. Cell 157, (2014).

74. Okuyama, T., Kitamura, T., Roy, D. S., Itohara, S. & Tonegawa, S. Ventral CA1 neurons store social memory. Science (80-.). 353, (2016).

75. Hitti, F. L. & Siegelbaum, S. A. The hippocampal CA2 region is essential for social memory. Nature 508, (2014).

76. Remedios, R. et al. Social behaviour shapes hypothalamic neural ensemble representations of conspecific sex. Nature 550, 388–392 (2017).

77. Li, Y. et al. Neuronal Representation of Social Information in the Medial Amygdala of Awake Behaving Mice. Cell 171, 1176–1190.e17 (2017).

78. Varea, O. et al. Synaptic abnormalities and cytoplasmic glutamate receptor aggregates in contactin associated protein-like 2/Caspr2 knockout neurons. Proc. Natl. Acad. Sci. U. S. A. 112, (2015).

79. Anderson, G. R. et al. Candidate autism gene screen identifies critical role for cell-adhesion molecule CASPR2 in dendritic arborization and spine development. Proc. Natl. Acad. Sci. U. S. A. 109, 18120–5 (2012).

80. Scott, R. et al. Loss of Cntnap2 Causes Axonal Excitability Deficits, Developmental Delay in Cortical Myelination, and Abnormal Stereotyped Motor Behavior. Cereb. Cortex (2017). doi:10.1093/cercor/bhx341

81. Selimbeyoglu, A. et al. Modulation of prefrontal cortex excitation/inhibition balance rescues social behavior in CNTNAP2 –deficient mice. Sci. Transl. Med. 9, eaah6733 (2017).

82. Behrmann, M. & Minshew, N. J. Sensory Processing in Autism. 180, 54–67 (2015).

83. Weinger, P. M., Zemon, V., Soorya, L. & Gordon, J. Low-contrast response deficits and increased neural noise in children with autism spectrum disorder. Neuropsychologia 63, 10–18 (2014).

